# The entorhinal cortex modulates trace fear memory formation and neuroplasticity in the lateral amygdala via cholecystokinin

**DOI:** 10.1101/2021.04.18.440346

**Authors:** Hemin Feng, Junfeng Su, Wei Fang, Xi Chen, Jufang He

## Abstract

Although the neural circuitry underlying fear memory formation is important in fear-related mental disorders, it is incompletely understood. Here, we utilized trace fear conditioning to study the formation of trace fear memory. We identified the entorhinal cortex (EC) as a critical component of sensory signaling to the amygdala. Moreover, we used the loss of function and rescue experiments to demonstrate that release of the neuropeptide cholecystokinin (CCK) from the EC is required for trace fear memory formation. We discovered that CCK-positive neurons extend from the EC to the lateral nuclei of the amygdala (LA), and inhibition of CCK-dependent signaling in the EC prevented long-term potentiation of sensory signals to the LA and formation of trace fear memory. Altogether, we suggest a model where sensory stimuli trigger the release of CCK from EC neurons, which potentiates sensory signals to the LA, ultimately influencing neural plasticity and trace fear memory formation.

## Introduction

Learning to associate environmental cues with subsequent adverse events is an important survival skill. Pavlovian fear conditioning is widely used to study this association and is performed by pairing a neutral stimulus (conditioned stimulus, CS), such as a tone, with a punishing stimulus (unconditioned stimulus, US), such as a shock (Pavlov, 1927). The CS-US pair elicits fear behaviors, including freezing and fleeing, which are often species-specific. Canonical delay fear conditioning is performed by terminating the CS and US at the same time. However, conditioned and unconditioned stimuli do not necessarily occur simultaneously in nature, and the brain has evolved mechanisms to associate temporally distinct events. Trace fear conditioning is used to study these mechanisms by inserting a trace interval between the end of the CS and the beginning of the US. The temporal separation between the CS and the US substantially increases the difficulty of learning as well as the recruitment of brain structures (Crestani et al., 2002; Runyan et al., 2004). Although trace fear conditioning provides essential insight into the neurobiology of learning and memory, many unanswered questions remain. For instance, the detailed neural circuitry underlying the formation of this trace fear memory and the potential modulatory chemicals involved in this process need to be further characterized.

Synaptic plasticity is the basis of learning and memory and refers to the ability of neural connections to become stronger or weaker. Long-term potentiation (LTP) is one of the most widely-studied forms of synaptic plasticity. The lateral nucleus of the amygdala (LA) receives multi-modal sensory inputs from the cortex and thalamus and relays them into the central nucleus of the amygdala (CeA), which then innervates the downstream effector structures (Phelps & LeDoux, 2005). LTP is developed in the auditory input pathway that signals to the LA. Auditory-responsive units in the LA fire faster after auditory-cued fear conditioning (Quirk et al., 1995). Optogenetic manipulation of the auditory input terminals in the LA leads to the suppression or recovery of LTP in the LA and can correspondingly suppress or recover conditioned fear responses (Nabavi et al., 2014). Together, these studies demonstrate that synaptic plasticity in the LA is impressively correlated with the formation of fear memory.

In addition to the amygdala, other neural regions, including the hippocampus (Bangasser, 2006), anterior cingulate cortex (ACC) (Han et al., 2003), medial prefrontal cortex (mPFC) (Runyan et al., 2004), and entorhinal cortex (EC) (Ryou et al., 2001), take part in trace fear conditioning.

The EC is integrated in the spatial and navigation systems of the animal (Fyhn et al., 2004; Hafting et al., 2005) and is essential for context-related fear associative memory (Maren & Fanselow, 1997). Moreover, the EC functions as a working memory buffer in the brain to hold information for temporal associations (Fransén, 2005; Schon et al., 2016). Here, a scenario of the dependence on the EC to associate the temporally-separated CS and US is manifested.

The neuropeptide cholecystokinin (CCK) is universally accepted as the most abundant neuropeptide in the central nervous system (CNS) (Rehfeld, 1978). CCK is recognized by two receptors in the CNS: CCK A receptor (CCKAR) and CCK B receptor (CCKBR). Previous studies in our laboratory unveiled that CCK and CCKBR enabled neuroplasticity as well as associative memory between two sound stimuli and between visual and auditory stimuli in the auditory cortex (X. Chen et al., 2019; Li et al., 2014; Z. Zhang et al., 2020). CCK and its receptors are intrinsically involved in fear-related mental disorders including anxiety (Q. Chen et al., 2006), depression (Shen et al., 2019), and post-traumatic stress disorder (PTSD) (Joseph et al., 2013). Moreover, the CCKBR agonist CCK-tetrapeptide (CCK-4) induces acute panic attacks in individuals with a panic disorder as well as in healthy human subjects (Bradwejn, 1993). Despite the clear connection between CCK and fear-related disorders, it remains elusive that the involvement of CCK in Pavlovian fear conditioning and the formation of cue-specific fear memory, which is possibly the neural foundation of these disorders.

In the present study, we investigated the involvement of CCK-expressing neurons in the EC in trace fear memory formation. We then examined how CCK enabled neuroplasticity in the auditory pathway to the LA by conducting the *in vivo* recording in the LA. Finally, we studied the contribution of the pathway from the EC to LA on the formation of trace fear memory in a physiological and behavioral context.

## Results

### Loss of CCK results in deficient trace fear memory formation in CCK^-/-^mice

The first question we asked here was whether CCK is involved in trace fear memory formation. We studied transgenic CCK^-/-^ mice (Cck-CreER, strain #012710, Jackson Laboratory), which lack CCK expression (X. Chen et al., 2019). We subjected CCK^-/-^ and wildtype control (WT, C57BL/6) mice to trace fear conditioning using two training protocols: long trace interval and short trace interval training.

Trace fear conditioning was performed by collecting baseline readouts on pre-conditioning day, training with the appropriate CS-US pairings on conditioning days, and testing the conditioned fear responses on post-conditioning/testing day. In the long trace protocol, mice sequentially received a 10-second pure tone (as the CS), a 20-second gap (trace interval), and a 0.5-second foot shock (as the US) (Figure 1a). We calculated the percentage of time frames where mice displayed a freezing response as the measure of fear memory. Freezing percentages were compared before (baseline) and after (post-training) trace fear conditioning as well as before (Figure 1b) and after (Figure 1c) presentation of the CS. At baseline, CCK^-/-^ (N = 10) and WT (N = 14) mice showed similarly low freezing percentages both before (Figure 1b) and after (Figure 1c) the CS (Figure 1b, two-way repeated-measures analysis of variance [RM ANOVA], significant interaction, F [1,22] = 4.65, P < 0.05; pairwise comparison, WT vs. CCK^-/-^ before CS, 7.0% ± 1.1% vs. 5.9% ± 0.8%; Bonferroni test, *P* > 0.05; Figure 1c, two-way RM ANOVA], significant interaction, F [1,22] = 13.87, *P* < 0.05; pairwise comparison, WT vs. CCK^-/-^ after CS, 9.9% ± 1.6% vs. 9.6% ± 1.5%, Bonferroni test, *P* > 0.05). After conditioning, CCK^-/-^ mice showed significantly lower freezing percentages (32.5% ± 6.2%) than WT mice after receiving the CS (61.6% ± 4.6%, pairwise comparison, *P* < 0.01), indicating poor performance in associating the CS with the US (Figure 1c, Movie S1, S2). This effect was not due to elevated basal freezing levels caused by training in WT animals (Figure 1b). Instead, we found that CCK^-/-^ mice (20.7% ± 3.0%) had slightly higher freezing percentages than WT mice (14.0% ± 1.7%) in the absence of the CS (pairwise comparison, *P* > 0.05). Together, these results suggest that trace fear conditioning results in elevated conditioned freezing percentages in WT mice, which are primarily elicited by the CS, and that loss of CCK impairs the freezing response to the CS. Furthermore, we defined an empirical threshold of moving velocity and converted the moving velocity to a binary freezing score plot, in which value 1 represents active status, and value 0 represents freezing status (see <u>Methods</u>). Using this method, we were able to assess the freezing response of the animal as it occurred during the CS presentation. Again, we found that WT mice obtained higher average freezing scores than CCK^-/-^ mice during the presentation of the CS (Figure 1d, \**P* < 0.05, two-sample t-test).

**Figure 1.**
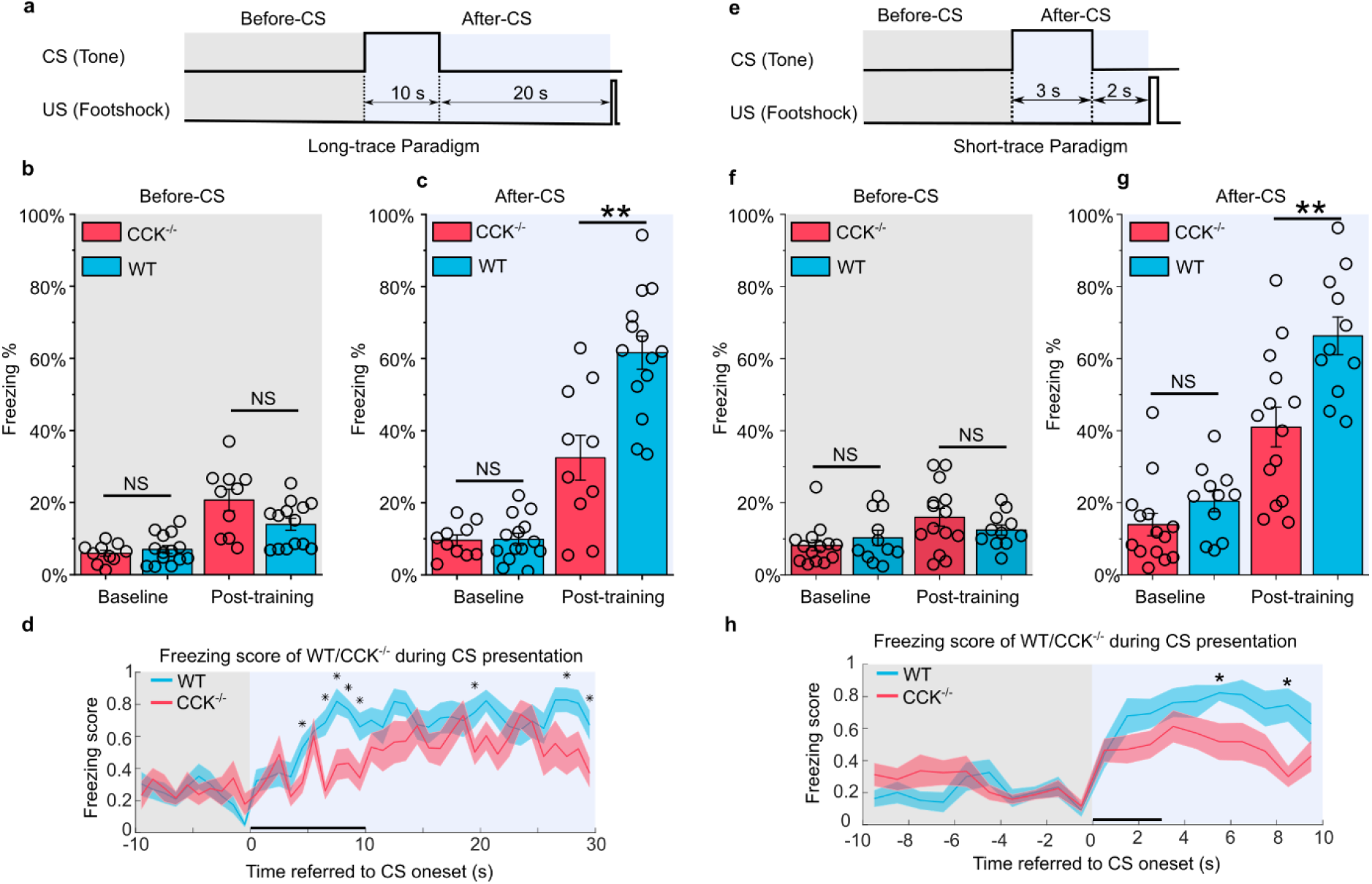
Trace fear memory formation deficit in CCK^-/-^mice. (**a**) Schematic diagram of the fear conditioning paradigm with a long trace interval of 20 s. Gray and light blue shadowed areas indicate the time frames before and after the onset of the CS (Before CS, After CS). CS, conditioned stimulus; US, unconditioned stimulus. (**b–c**) Freezing percentages before (b) and after (c) the CS. Freezing percentages were recorded at baseline on the pre-conditioning day and post-training on the post-conditioning day. WT, wildtype, N = 14; CCK^-/-^, CCK-knockout, N = 10. \**P* < 0.05; \*\**P* < 0.01; \*\*\**P* < 0.001; NS, not significant. Statistical significance was determined by two-way RM ANOVA with Bonferroni post-hoc pairwise test. RM ANOVA, repeated measures analysis of variance. (**d**) Freezing score plot of the two groups of mice during the testing session. Solid lines indicate the mean value, and shadowed areas indicate the SEM. The black bar indicates the presence of the CS from 0 s to 10 s. \**P* < 0.05; two-sample t-test; SEM, standard error of the mean. (**e**) Schematic diagram of the fear conditioning paradigm with a short trace interval of 2 s. (**f–g**) Freezing percentages before (**f**) and after (**g**) the CS. WT, N = 11; CCK^-/-^, N = 14. (**h**) Freezing score plot of the two groups of mice during the testing session. The black bar indicates the presence of the CS from 0 s to 3 s. \**P* < 0.05; two-sample t-test.

In addition to the long trace interval, we also investigated freezing responses of mice during a short trace fear conditioning paradigm. Mice were presented a 3-second CS followed by a 2-second trace interval and a 0.5-second electrical foot shock (Figure 1e). Before training, WT (N = 11) and CCK^-/-^ (N = 14) mice showed similarly low freezing percentages both before (Figure 1f) and after (Figure 1g) presentation of the CS (Figure 1g, two-way RM ANOVA, significant interaction, F [1,23] = 4.85, *P* < 0.05; pairwise comparison, WT vs. CCK^-/-^ in the baseline session, 20.4% ± 2.9% vs. 13.9% ± 3.1%, *P* > 0.05; Figure 1f, two-way RM ANOVA, interaction not significant, F [1,23] = 1.8, *P* = 0.19 > 0.05; pairwise comparison, WT vs. CCK^-/-^ in the baseline session, 10.3% ± 2.1% vs. 8.2% ± 1.5%, *P* > 0.05). Consistent with results from the long trace paradigm, CCK^-/-^ mice showed an impaired freezing response (41.0% ± 5.5%) to the CS after training compared to WT mice (66.3% ± 5.2%, pairwise comparison, *P* < 0.01, Figure 1g, <u>Movie S3, S4</u>). Additionally, we observed no significant difference between fear conditioned WT and CCK^-/-^ mice prior to the presentation of the CS (Figure 1f, pairwise comparison, WT vs. CCK^-/-^ in the post-training session, 12.4% ± 1.4% vs. 16.0% ± 2.4%, *P* > 0.05). Finally, we found significant differences in freezing scores between WT and CCK^-/-^ mice when presented the CS (Figure 1h, \**P* < 0.05, two-sample t-test).

We conducted the innate hearing and fear expression examinations to rule out a potential inherent deficit derived from genome editing in CCK^-/-^ transgenic mice. To evaluate hearing, we recorded the open-field auditory brainstem response (ABR) in anesthetized animals. We observed five peaks in both WT and CCK^-/-^ mice at sound intensities above 50 dB of sound pressure level (dB SPL) (Figure S1b), and we did not observe any remarkable differences between the waveforms. Compared to WT mice, CCK^-/-^ mice had better hearing (40.0 ± 1.2 dB in CCK^-/-^ mice, N = 15, vs. 47.3 ± 2.1 dB in WT mice, N = 11, two-sample t-test, *P* < 0.01, Figure S1c). Thus, auditory perception does not account for the deficient trace fear memory formation of CCK^-/-^ mice.

As fear expression is the behavioral output of fear conditioning, we wondered if CCK^-/-^ mice suffered from a deficit in fear expression, which is observed in Klüver-Bucy syndrome and other diseases (Lilly et al., 1983). To test whether the CCK^-/-^ mice have a deficit in fear expression, we presented a loud (90 dB SPL) white noise and quantified sound-driven innate freezing. We found no statistical difference between WT (46.1% ± 5.5%, N = 11) and CCK^-/-^ mice (46.5% ± 6.6%, N = 14, two-sample t-Test, *P* > 0.05, Figure S1d), indicating that CCK^-/-^ mice can express passive defensive behaviors such as freezing. Thus, the deficient trace fear memory formation of CCK^-/-^ is not due to a deficit in fear expression and may be due to a deficit in establishing an association between the CS and the US.

In summary, CCK^-/-^ mice display deficient trace fear memory formations in both short and long trace models that are not caused by inherent abnormalities in hearing or fear expression.

### Deficient neural plasticity in the LA of CCK^-/-^mice

As neural plasticity in the LA is widely regarded as the basis of fear memory formation (Kim & Cho, 2017; LeDoux, 2000; Nabavi et al., 2014; Rogan et al., 1997), we examined LTP in the LA of WT and CCK^-/-^ mice by *in vivo* recording (Figure 2a). First, we successfully recorded the auditory evoked potential (AEP) in the LA of anesthetized WT and CCK^-/-^ mice (Figure 2b–e). Then, we used theta-burst electrical stimulation (TBS) to induce LTP of AEP (AEP-LTP) (Figure 2f). Interestingly, AEP-LTP was effectively induced in WT mice (N = 15) but was not in CCK^-/-^ mice (N = 12). WT mice demonstrated remarkable potentiation (Figure 2g, two-way RM ANOVA, significant interaction, F [1,25] = 6.8, *P* = 0.015 < 0.05; pairwise comparison, after vs. before induction, 142.7% ± 17.5% vs. 99.1% ± 2.8%, *P* = 0.011 < 0.05), whereas CCK^-/-^ mice showed no potentiation (pairwise comparison, after vs. before induction, 98.0% ± 5.8% vs. 100.6% ± 3.4%, *P* > 0.05). These results suggest that CCK^-/-^ mice have a deficit in neural plasticity in the LA that may contribute to their reduced response to trace fear conditioning.

**Figure 2.**
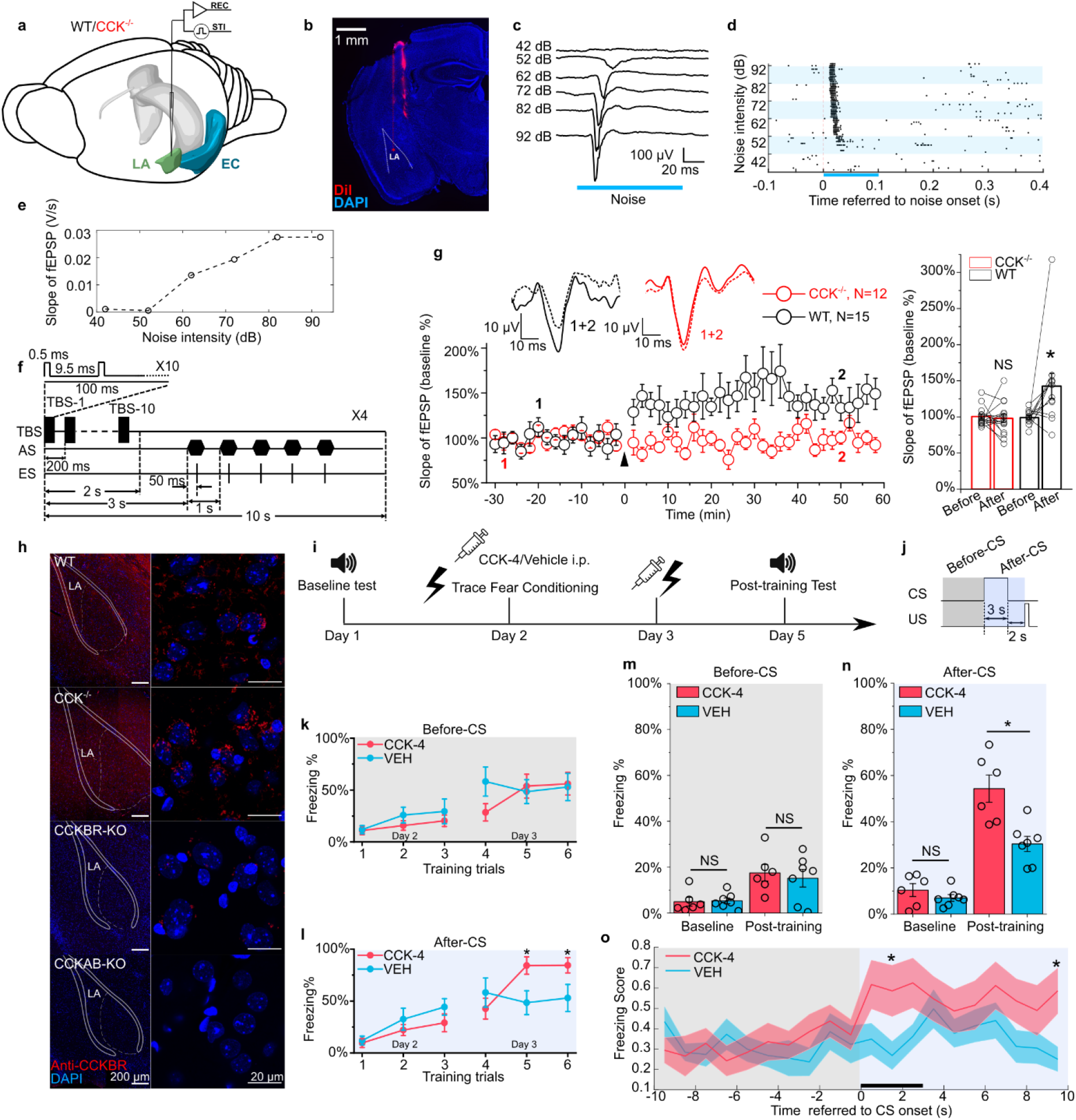
Neural plasticity deficit in the LA of CCK^-/-^mice and the rescuing effect of exogenous CCK. (**a**) Schematic diagram of *in vivo* recording in the LA. EC, entorhinal cortex; LA, lateral amygdala. STI, stimulation. REC, recording. (**b**) Post-hoc verification of electrode tracks and recording area. (**c**) Representative AEP traces in response to different levels of noise stimulus. AEP, auditory evoked potential. (**d**) Representative traces of multiunit spikes to different levels of noise stimulus. (**e**) Representative input/output (I/O) curve of the slope of AEP versus noise intensity. fEPSP, field excitatory postsynaptic potential. (**f**) Schematic diagram of the pairing protocol to induce LTP of AEP via theta-burst stimulation (TBS). LTP, long term potentiation; ES, electrical stimulation; AS, auditory stimulation. (**g**) Time course plot of the normalized AEP slope during LTP. The WT group is indicated in black, and the CCK^-/-^ group is indicated in red. Representative traces of the AEP before (dotted line) and after (solid line) TBS are shown in inset panels for both groups. The average normalized slopes 10 min before pairing (–10–0 min, before) and 10 min after pairing (50–60 min, after) in the two groups of mice are shown on the right. \*\*\**P* < 0.05; two-way RM ANOVA with Bonferroni post-hoc pairwise test; RM ANOVA, repeated measures analysis of variance; NS, not significant. (**h**) Immunofluorescent staining of CCK B receptor (CCKBR) in brain slices from WT, CCK^-/-^, CCKBR-KO, and CCKAB-KO mice. Magnified images are shown on the right. CCKBR-KO, CCK B receptor knock-out mouse; CCKAB-KO, CCK A receptor and B receptor double knock-out mouse. (**i**) Experimental timeline for (**j–o**). (**j**) Schematic diagram of the CS-US presentation. Gray and light blue shadowed areas indicate the time frames before and after CS presentation (Before-CS, After-CS). (**k–l**) Freezing percentages before (**k**) and after (**l**) the CS during fear conditioning training on training day. Animals underwent six trials during a 2-day training (day 2 and 3). CCK-4, N = 6; VEH, N = 6; \**P* < 0.05; two-sample t-test. (**m–n**) Freezing percentages before (**m**) and after (**n**) the CS on the pre-training day (baseline) and the post-training day. CCK-4, N = 6; VEH, N = 6; \**P* < 0.05; NS, not significant; two-way RM ANOVA with Bonferroni post-hoc pairwise test; RM ANOVA, repeated measures analysis of variance. (**o**) Freezing score plot of the two groups of mice during the testing session on day 5. Solid lines indicate the mean value, and shadowed areas indicate the SEM. The black bar indicates the presence of the CS from 0 s to 3 s. \**P* < 0.05; two-sample t-test; SEM, standard error of the mean.

### Stimulation of CCKBR rescues the formation of trace fear memory in CCK^-/-^mice

Although the translation and release of CCK are disrupted in CCK^-/-^ mice, we found that the predominant CCK receptor, CCKBR, was expressed normally in both WT and CCK^-/-^ mice (Figure 2h). Therefore, we hypothesized that exogenous stimulation of CCKBR might rescue trace fear memory deficits in CCK^-/-^ mice. CCKBR can be stimulated by several agonists, including CCK octapeptide sulfated (CCK-8s) and CCK tetrapeptide (CCK-4). As CCK-8s is a potent agonist of both CCKAR and CCKBR, we selected CCK-4, which is a preferred CCKBR agonist (Berna et al., 2007). To monitor CCK signaling *in vivo*, we expressed a G protein-coupled receptor (GPCR)-activation-based CCK sensor (GRAB_CCK_, AAV-hSyn-CCK2.0) in the LA of CCK^-/-^ mice (Jing et al., 2019). Using this model, binding of the GPCR CCKBR with endogenous or exogenous CCK results in increased fluorescence intensity, which we measured by fiber photometry in the LA (Figure S2a). We first confirmed that intraperitoneal (i.p.) administration of CCK-4 permeated the blood-brain-barrier (BBB) and activated the CCK2.0 sensor. Moreover, we demonstrated that administration of CCK-4 evoked a clear and long-term increase in the fluorescent signal (Figure S2b). Together, these data verify that CCK-4 passes through the BBB and binds with CCKBR in the LA.

After validating our model, we conducted short trace fear conditioning in CCK^-/-^ mice on two consecutive days just after intraperitoneal administration of CCK-4 or the corresponding vehicle (VEH) (Figure 2i–j). We collected data during the two conditioning days to monitor the learning curve of mice as conditioning progressed. The learning curves were plotted as the freezing percentages of CCK-4- or VEH-treated CCK^-/-^ mice during the six training trials (Figure 2k–l). During the first three trials on the first conditioning day and even in the fourth trial on the second conditioning day, we did not observe any statistical differences between the two groups. During the fifth and sixth training trials conducted on the second conditioning day, we found that CCK-4-treated mice had significantly higher freezing levels than VEH-treated mice (Figure 2l, 84.2% ± 8.4% in the CCK-4 group [N = 6] vs. 48.4% ± 11.5% in the VEH group [N = 7] in the fifth trial; 84.4% ± 7.3% in the CCK-4 group vs. 52.9% ± 13.0% in the VEH group in the sixth trial, two-sample t-test, both *P* < 0.05). In support of this evidence, we did not find a statistical difference between the two groups prior to CS presentation during the fifth or sixth trials (Figure 2k, 53.8% ± 11.5% in the CCK-4 group vs. 52.5% ± 11.8% in the VEH group in the fifth trial; 56.0% ± 10.8% in the CCK-4 group vs. 47.8% ± 11.8% in the VEH group in the sixth trial, two-sample t-test, both *P* > 0.05). Together, these data suggest that mice in the CCK-4- and VEH-treated groups showed similar baseline freezing levels and that CCK-4 treatment improved trace fear conditioning learning responses in CCK^-/-^ mice.

We went on to assess the conditioned fear response in CCK-4- and VEH-treated CCK^-/-^ mice two days after training in comparison to fear responses at baseline prior to training (Figure 2m–n). We found that CCK4-treated mice showed remarkably higher freezing levels than VEH-treated mice post-training, whereas no significant difference was detected at baseline (Figure 2n, two-way RM ANOVA, significant interaction, F [1,11] = 6.40, *P* = 0.028 < 0.05; pairwise comparison, CCK-4 vs. VEH at baseline, 10.4% ± 2.8% vs. 7.0% ± 1.4%, *P* > 0.05; CCK-4 vs. VEH post-training, 54.3% ± 5.9% vs. 30.4% ± 3.3%, *P* < 0.05; <u>Movie S5, S6</u>). There was no statistical difference between the two groups before the presentation of the CS (Figure 2m, two-way RM ANOVA, the main effect of drug application [CCK-4 vs. VEH] on freezing percentage was not significant, F [1,11] = 0.15, *P* = 0.70). Additionally, CCK-4-treated mice had significantly higher freezing scores than VEH-treated mice (Figure 2o). These results indicate that CCK-4 treatment effectively improved learning response to trace fear conditioning in CCK^-/-^ mice. Moreover, this rescue was not an artifact caused by reduced locomotion after drug application and fear conditioning training, as there was no difference between the two groups in the freezing percentage prior to presentation of the CS (Figure 2m). Therefore, the exogenous application of a CCKBR agonist activated endogenous CCKBR and improved the fear memory formation of CCK^-/-^ mice after trace fear conditioning.

### CCK neurons in the EC are critical for the formation of the trace fear memory

We next examined the source of endogenous CCK that signals to the LA. We injected a potent retrograde neuronal tracer Cholera Toxin Subunit B (CTB) conjugated to a fluorescent tag Alexa-647 (CTB-647) into the LA and dissected the upstream anatomical brain regions that project to the LA (Figure 3a). In addition to regions that are canonically involved in fear circuitry, including the auditory cortex (AC) and the medial geniculate body (MGB), we found that EC was also densely labeled with retrograde CTB-647, suggesting that the EC is connected with the LA (Figure 3b-e). We next injected a Cre-dependent retrograde AAV (retroAAV-hSyn-FLEX-jGcamp7s) into the LA of CCK-ires-Cre (CCK-Cre) mice to label CCK-positive neurons that project into the LA, to further confirm the above observation (Figure 3f-g). In CCK-ires-Cre mouse line, Cre expression was restricted to the CCK-expressing neurons, where the Cre-mediated recombination took place and the Cre-dependent green, fluorescent protein jGcamp7s was expressed (Figure 3f). Fluorescent signal was detected in the AC and the EC, but not in the MGB (Figure 3h–j), which suggests that CCK may originate from these two brain regions during trace fear memory formation. Immunofluorescent staining revealed that most CCK-positive neurons in the EC that project to the LA are glutamatergic (Figure 3k–l), which is consistent with our previous findings in CCK-positive neurons in the EC (X. Chen et al., 2019).

**Figure 3.**
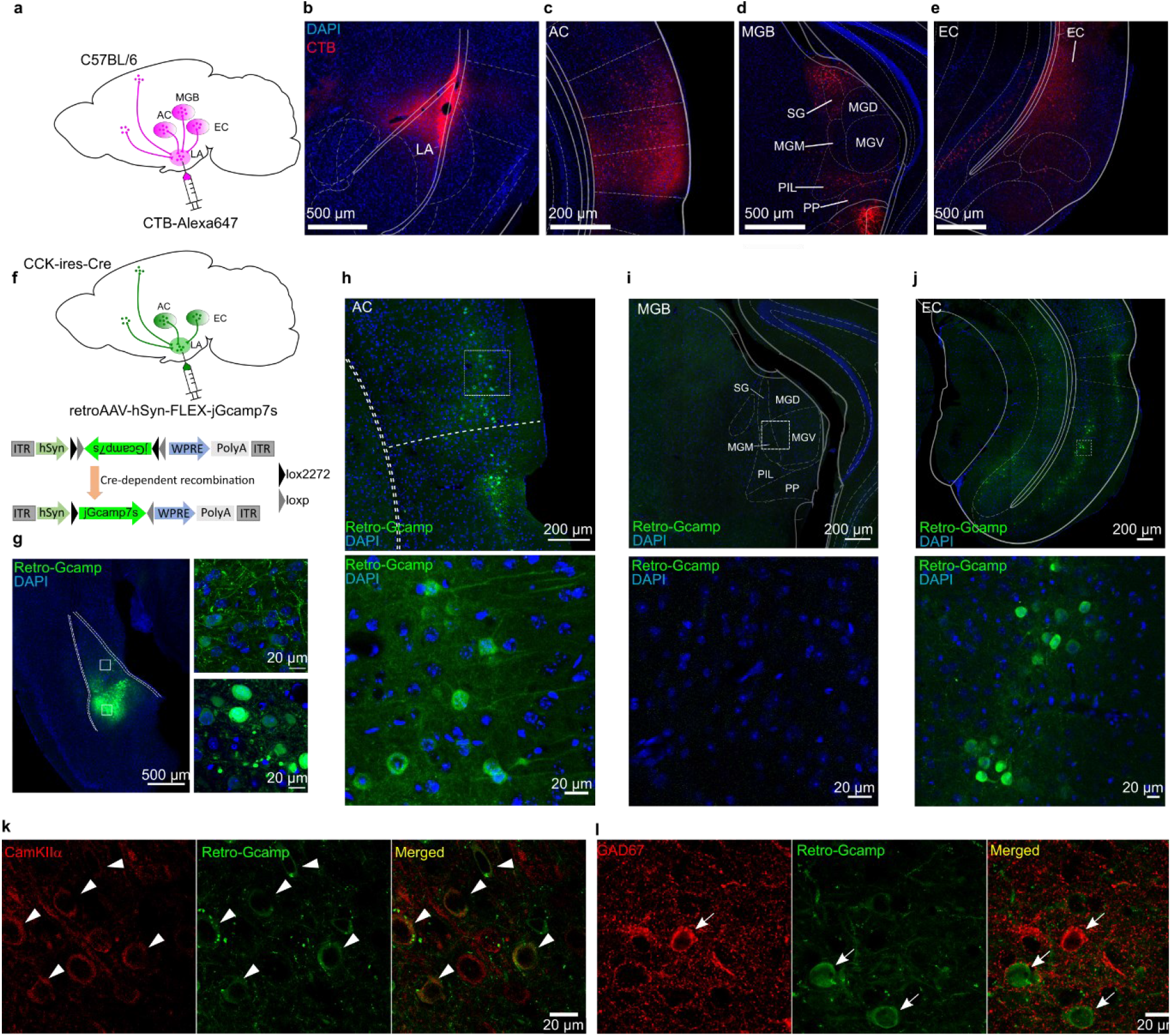
Dissection of inputs of the LA with retrograde tracer and virus. (**a**) Schematic diagram of retrograde tracing with Alexa647-conjugated cholera toxin subunit B (CTB). (**b–e**) Representative fluorescent images of the injection site of the CTB tracer (**b**), the canonical upstream regions, including the auditory cortex (**c**) and the auditory thalamus (**d**), and the non-canonical entorhinal cortex (**e**). AC, auditory cortex; MGB, medial geniculate body; SG, suprageniculate thalamic nucleus; MGM, medial MGB; PIL, posterior intralaminar thalamic nucleus; PP, peripeduncular nucleus; EC, entorhinal cortex. (**f**) Schematic diagram of cell type-specific retrograde tracing with Cre-dependent retrograde AAV (retroAAV-hSyn-FLEX-jGcamp7s). (**g**) Verification of the injection site in the LA. Magnified images are shown in insets on the right. Retro-Gcamp, retrograde jGcamp7s signal. (**h–j**) Retrograde signals in the AC (**h**), MGB (**i**), and EC (**j**). Magnified images are shown in the bottom insets. (**k–l**) Co-immunofluorescent staining of retrograde tracing of the LA with either the excitatory neuronal marker CamKIIα (**k**) or the inhibitory neuronal marker GAD67 (**l**).

Interestingly, the EC is involved in the formation of trace fear memory but is not a component of canonical delay fear memory (Esclassan et al., 2009). This selectivity suggests that the EC may be a component of the neural circuit underlying trace fear memory formation. To evaluate a requirement for the EC in trace fear memory, we utilized a Designer Receptors Exclusively Activated by Designer Drugs (DREADD) system to silence EC neurons (Armbruster et al., 2007). Specifically, the inhibitory receptor hM4Di was expressed in the EC of WT mice (Figure 4a) and was activated by administration of the designer drug clozapine (CLZ). Activation of hM4Di by CLZ induces membrane hyperpolarization, effectively silencing neurons. We verified EC neuron silencing by *in vivo* electrophysiological recording (Figure 4b–d and Figure S3). We found that a low dose of CLZ (0.5 mg/kg) effectively suppressed both instant and long-term neuronal firing. Of note, we used CLZ instead of the canonical DREADD ligand clozapine-N-oxide (CNO). A recent study identified CLZ as the active metabolite of CNO (Gomez et al., 2017), and CLZ more effectively penetrates the BBB and binds with DREADD receptors compared to CNO. As a result, a much lower dose of CLZ can elicit similar behavioral effects as higher doses of CNO (Gomez et al., 2017). Therefore, we used a low dose of CLZ (0.5 mg/kg) in our experiments.

**Figure 4.**
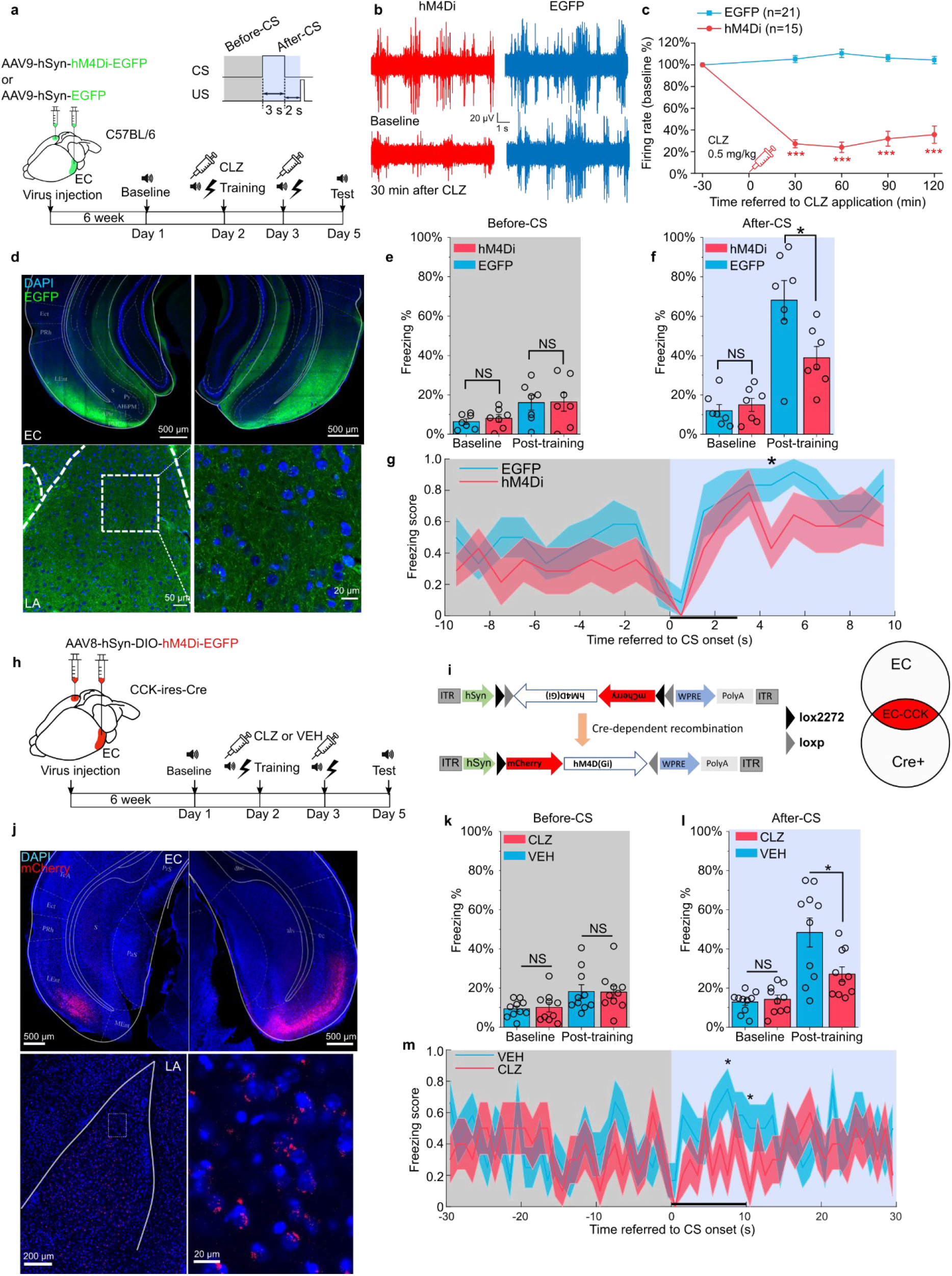
Formation of trace fear memory is suppressed by chemogenetic inhibition of the EC and CCK-positive EC neurons. (**a**) Schematic diagram of trace fear conditioning and chemogenetic inhibition of the EC. EC, entorhinal cortex; hM4Di, inhibitory DREADD receptor; CLZ, clozapine. (**b**) Representative traces of extracellular recording in the EC before and after systemic application of CLZ in hM4Di-expressing (red) and EGFP-expressing mice (blue). (**c**) Normalized firing rate of the EC neurons before and after systemic CLZ application. ***P < 0.001; two-sample t-test. (**d**) Verification of viral expression in the bilateral EC (top panel) and the EC-LA projection (bottom left panel). A magnified image of the EC-LA projection is shown in the bottom right inset. (**e–f**) Freezing percentages before (e) and after (f) the CS during the testing session in hM4Di-expressing (N = 7) or EGFP-expressing mice (N = 7). *P < 0.05; NS, not significant; two-way RM ANOVA with Bonferroni post-hoc pairwise test; RM ANOVA, repeated measures analysis of variance. (**g**) Freezing score plot of hM4Di-expressing and EGFP-expressing mice during the testing session. Solid lines indicate the mean value, and shadowed areas indicate the SEM. The black bar indicates the presence of the CS from 0 s to 3 s. *P < 0.05; two-sample t-test; SEM, standard error of the mean. (**h–i**) Schematic diagrams of chemogenetic CCK inhibition in the EC. Cre-dependent hM4Di was expressed in CCK-Cre mice. After Cre-mediated recombination, CCK neurons in the EC were transfected with hM4Di. (**j**) Verification of viral expression in the bilateral EC (top panel) and the EC-LA projection (bottom left panel). A magnified image of the EC-LA projection is shown in the bottom right inset. (**k-l**) Freezing percentages before (**k**) and after (**l**) the CS during the testing session in mice treated with CLZ or vehicle (VEH). *P < 0.05; NS, not significant; two-way RM ANOVA with Bonferroni post-hoc pairwise test. (**m**) Freezing score plot of CLZ- and VEH-treated mice during the testing session. The black bar indicates the presence of the CS from 0 s to 10 s. *P < 0.05; two-sample t-test; SEM, standard error of the mean.

Six weeks after injection of AAV9-hSyn-hM4Di-EGFP or AAV9-hSyn-EGFP, we administered CLZ by intraperitoneal injection and conducted short trace fear conditioning 30 min later. We repeated the CLZ treatment and trace fear conditioning the following day and tested conditioned fear responses two days after that. As expected, mice expressing hM4Di (hM4Di, N = 7) showed significantly lower freezing percentages in response to the CS than those expressing the control virus (EGFP, N = 7) post-training (Figure 4f, two-way RM ANOVA, significant interaction, F [1,12] = 7.42, *P* = 0.018 < 0.05; EGFP vs. hM4Di post-training, 68.1% ± 10.0% vs. 39.0% ± 5.7%, *P* = 0.035 < 0.05; Movie S7, S8). No significant differences were observed between the two groups at baseline (Figure 4f, pairwise comparison, EGFP vs. hM4Di at baseline, 12.0% ± 3.1% vs. 15.0% ± 3.3%, *P* > 0.05) or prior to the CS (Figure 4e, two-way RM ANOVA, interaction not significant, F [1, 12] = 0.05, *P* = 0.82 > 0.05; pairwise comparison, EGFP vs. hM4Di post-training, 16.0% ± 3.8% vs. 16.4% ± 4.7%, *P* > 0.05).

As we have shown that CCK-positive neural projections extend from the EC to the LA, we transfected CCK-expressing neurons in the EC with a Cre-dependent hM4Di in CCK-Cre mice (Figure 4h–j). These mice received an i.p. injection of CLZ (N = 10) or VEH (N = 10) prior to long trace fear conditioning. After training, mice injected with CLZ showed significantly lower freezing percentages than those injected with the VEH, whereas no statistical differences were observed at baseline or prior to the CS (Figure 4l, two-way RM ANOVA, significant interaction, F [1,18] = 5.90, *P* = 0.026 < 0.05; pairwise comparison, CLZ vs. VEH at baseline, 12.9% ± 1.7% vs. 14.2% ± 2.2%, *P* > 0.05; CLZ vs. VEH post-training, 48.4% ± 7.4% vs. 27.1% ± 3.7%, *P* = 0.017 < 0.05; Figure 4k, two-way RM ANOVA, interaction not significant, F [1, 18] = 0.043, *P* = 0.84 > 0.05; pairwise comparison, CLZ vs VEH at baseline, 10.2% ± 2.4 vs. 9.4% ± 1.4%, *P* > 0.05; CLZ vs. VEH post-training, 18.0% ± 3.2% vs. 18.3% ± 3.4%, *P* > 0.05; Movie S9, S10). These results mirror those observed in CCK^-/-^ mice and suggest that trace fear memory formation relies on intact and functional CCK-positive neurons in the EC.

### CCK-positive neuronal projections are predominant in the EC-LA pathway

To further demonstrate that afferents to the amygdala originate from CCK-expressing neurons in the EC, we locally injected a Cre-dependent color-switching virus (AAV-CAG-DO-mCherry-DIO-EGFP) in the EC of CCK-Cre mice (N = 2; Figure 5a–b). With this combination, CCK-positive neurons express EGFP, and CCK-negative neurons express mCherry (Saunders et al., 2012). We found that EGFP+ (i.e., CCK+) neurons made up a slightly higher proportion of labeled neurons than mCherry+ (i.e., CCK–) neurons (Figure 5c–d, EGFP vs. mCherry, 58.9% ± 4.8% vs. 38.6% ± 5.0%, one-way RM ANOVA, Wilks’ Lambda = 0.58, F [1,6] = 4.34, *P* = 0.0822 > 0.05). Interestingly, we found that CCK+ neural projections from the EC to the LA were densely labeled with EGFP, whereas mCherry labeling of CCK– projections was dramatically weaker. Quantitative analysis revealed that the projection intensity of the EC^CCK+^→ LA was 3-fold higher than the EC^CCK-^→ LA (35.6% ± 9.5%). In other words, CCK-positive afferents constituted approximately 75% of total afferents from the EC to the LA (Figure 5e–f).

**Figure 5.**
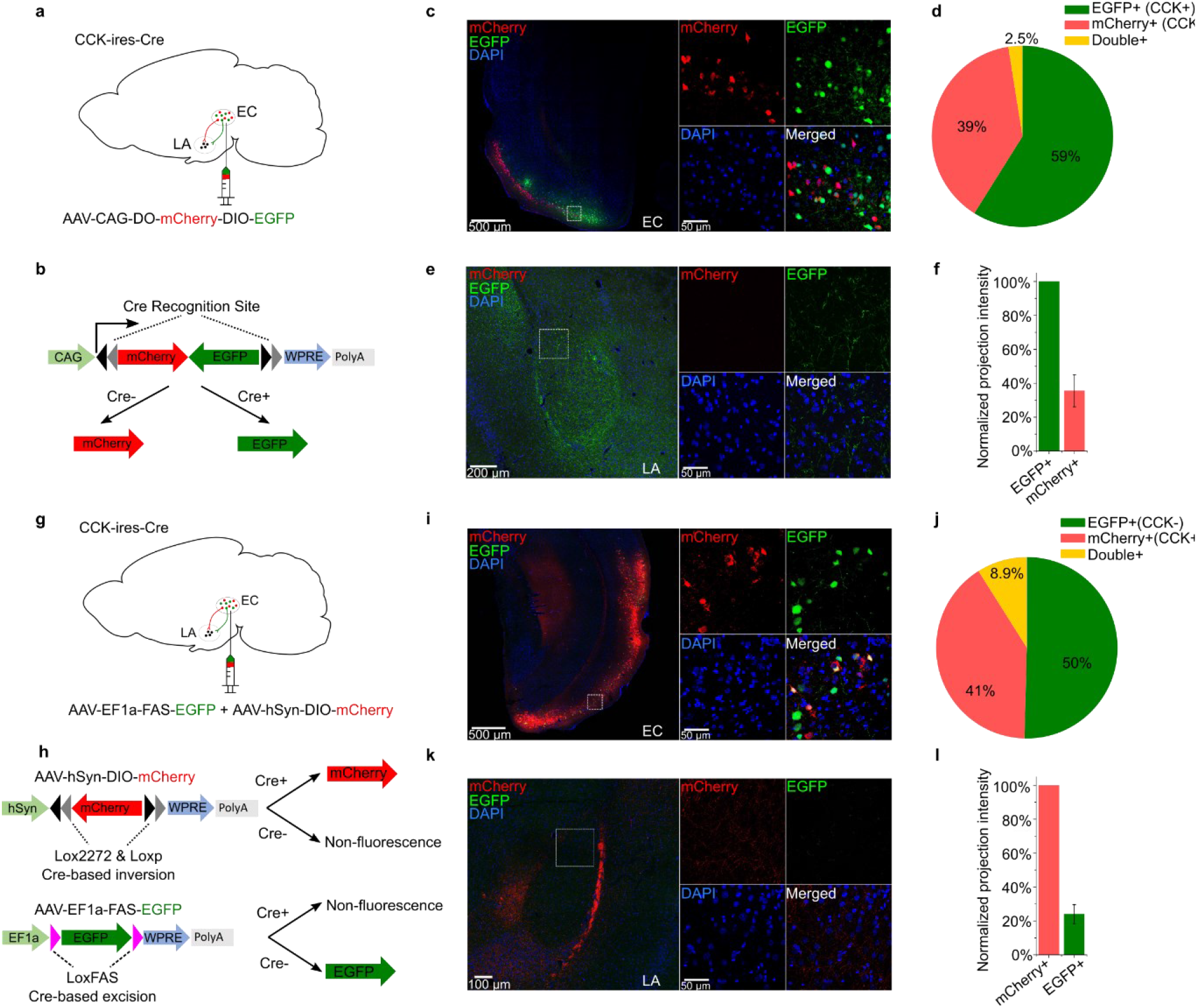
CCK-expressing projections predominate in the EC-LA pathway. (**a–b**) Schematic diagram of Cre-dependent color-switch labeling in the EC-LA pathway. AAV-CAG-DO-mCherry-DIO-EGFP was injected in the EC. Using this labeling scheme, EGFP is expressed in CCK+ neurons, and mCherry is expressed in CCK– neurons. (**c–d**) Visualization (**c**) and quantification (**d**) of viral expression in the EC. Representative immunofluorescent images in the EC 7 weeks after viral injection (**c**). Scale bar = 500 μm (left). Magnified images are shown in insets on the right. Scale bar = 50 μm. Percentages of EGFP+ (CCK+), mCherry+ (CCK–), and double-positive neurons (**d**). No statistical differences were observed. *P* = 0.08; one-way RM ANOVA, repeated measures analysis of variance. (**e–f**) Visualization (**e**) and quantification (**f**) of EGFP-expressing (CCK+) and mCherry-expressing (CCK–) afferents in the amygdala stemming from the EC. The fluorescent intensity of neuronal projections was normalized to the EGFP+ signal, which was approximately 3-fold stronger than the mCherry+ signal (35.6% ± 9.5%). (**g–h**) Schematic diagram of Cre-dependent color-switch labeling in the EC-LA pathway. A mixture of AAV-hSyn-DIO-mCherry and AAV-EF1α-FAS-EGFP was injected in the EC. Using this labeling scheme, mCherry is expressed in CCK+ neurons, and EGFP is expressed in CCK– neurons. (**i–j**) Visualization (**i**) and quantification (**j**) of viral expression in the EC. Representative immunofluorescent images in the EC 7 weeks after viral injection (**c**). Scale bar = 500 μm (left). Magnified images are shown in insets on the right. Scale bar = 50 μm. Percentages of mCherry+ (CCK+), EGFP+ (CCK-), and double-positive neurons (**j**). No statistical differences were observed. *P* = 0.55; one-way RM ANOVA; Wilks’ Lambda = 0.94; F (1,6) = 0.39. (**k–l**) Visualization (**k**) and quantification (**l**) of EGFP-expressing (CCK+) and mCherry-expressing (CCK–) afferents in the amygdala stemming from the EC. The fluorescent intensity of neuronal projections was normalized to the mCherry+ signal, which was approximately 4-fold stronger than the EGFP+ signal (24.0% ± 5.6%).

To determine if the fluorescent reporter proteins interfered with projection strength, we inverted the color combination by combining two AAVs: AAV-hSyn-DIO-mCherry and AAV-EF1α-FAS-EGFP (Saunders et al., 2012). These Cre-dependent AAVs were injected into the EC of CCK-Cre mice. In CCK-Cre mice, AAV-hSyn-DIO-mCherry induces Cre-ON mCherry expression in CCK+ neurons, and AAV-EF1α-FAS-EGFP induces Cre-OFF EGFP expression in CCK– neurons (Figure 5g–h). With the mixed AAVs, we labeled approximately 50% CCK– EGFP+ neurons, 41% CCK+ mCherry+ neurons, and 8.9% double-positive neurons (Figure 5i–j). The higher percentage of double-positive neurons present in this system indicates a higher probability of off-target effects compared to the previous color-switching AAV (8.9% ± 2.7% vs. 2.5% ± 1.1%). Consistent with the previous color-switching AAV, we observed that CCK+ (mCherry+) projections were predominant. Specifically, the intensity of the EC^CCK+^→ LA was approximately 4-fold higher than the EC^CCK-^→ LA (24.0% ± 5.6%). Altogether, our results suggest that the EC^CCK+^→ LA is the predominant subpopulation of projections, and that these projections are of functional significance in the EC-LA pathway.

### CCK-positive neural projections from the EC to the LA enable neural plasticity and modulate trace fear memory formation

Finally, we asked whether CCK-positive projections from the EC modulate neural plasticity in the LA. First, we expressed a Cre-dependent high frequency-responsive channelrhodopsin (ChR2) variant E123T (ChETA) under control of the universal EF1α promoter in CCK-Cre mice (Figure 6a). Then, we implanted optic fibers targeting the LA to illuminate EC^CCK+^→ LA projections and electrodes to conduct *in vivo* electrophysiological recording as before (Figure 6b). Post-hoc anatomical analysis confirmed the distribution of ChETA in the EC-LA axon terminals (Figure 6c). These CCK+ projections were innervated with postsynaptic CCKBR (Figure 6d), suggesting that CCK+ projections from the EC effectively activated CCKBR in the LA. Finally, we recorded auditory evoked potential (AEP) and visual evoked potential (VEP) in the LA of anesthetized mice (Figure 6e–g). Although AEP and VEP had similar waveforms, the latency of AEP was much shorter than VEP (Figure 6e–f, peak latency: 38.9 ± ms for AEP, N = 13, vs. 89.5 ± 3.1 ms for VEP, N = 11, two-sample t-test, *P* < 0.001). This observation implies that input pathways other than the canonical thalamo-cortico-amygdala and thalamo-amygdala projections regulate the transmission of visual cues. We applied high-frequency-laser-stimulation (HFLS, Figure 6h) of the EC-LA axons before the auditory stimulus (AS) to trigger AEP-LTP in the LA. After induction, the AEP slope in the ChETA-expressing group (n = 10) increased significantly, whereas the VEP slope did not change (Figure 6i–j, two-way RM ANOVA, significant interaction, F [1,9] = 14.46, *P* = 0.0042 < 0.01; pairwise comparison, AEP before vs. after pairing, 97.8% ± 5.5% vs. 187.6% ± 15.6%, *P* < 0.001; VEP before vs. after pairing, 96.3% ± 4.9% vs. 120.7% ± 9.1%, *P* = 0.67). Additionally, we injected a non-opsin expressing control AAV (AAV-EF1α-DIO-EYFP, n = 20) and the AEP-LTP was not induced with the same protocol (two-way RM ANOVA between CHETA and EYFP, F [1,30] = 46.65, *P* <0.001; pairwise comparison, before vs. after pairing in the EYFP group, 102.8% ± 2.2% vs. 106.7% ± 4.8%, *P* > 0.05, Figure 6h-i) These results suggest that high frequency activation of EC^CCK+^→ LA switches the AEP-LTP in the LA.

**Figure 6.**
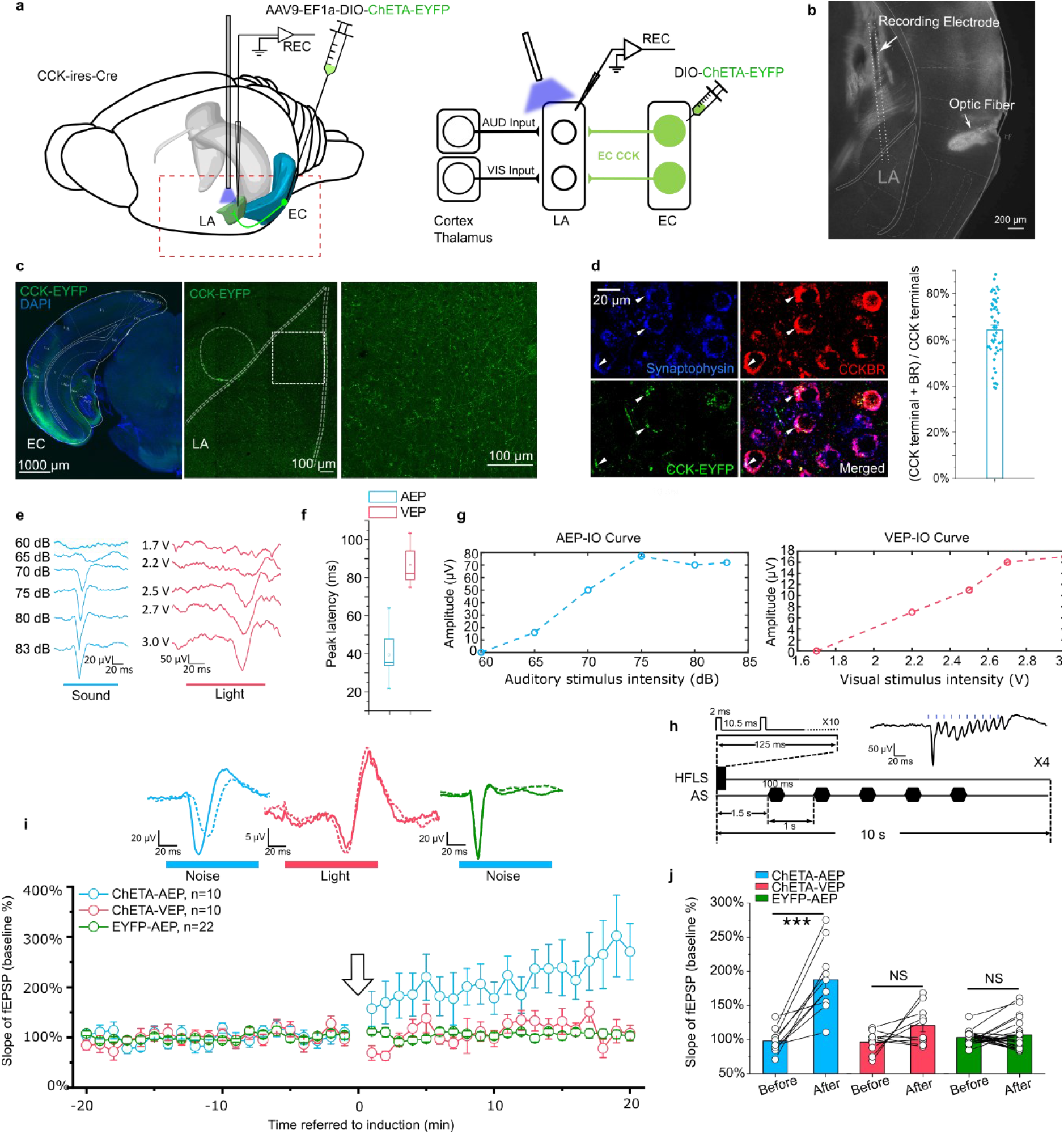
High frequency activation of the EC^CCK+^→ LA pathway induces LTP of AEP in the LA. (**a**) Schematic diagram of the experiment. The Cre-dependent high frequency-responsive opsin ChETA was expressed in the EC of CCK-Cre mice. Electrodes were inserted into the LA, and blue light was used to illuminate the recording area. The red rectangle in the left panel is magnified in the right panel to illustrate the neural pathways that are recruited during recording. AUD, auditory stimulus; VIS, visual stimulus; LA, lateral amygdala; EC, entorhinal cortex; REC, recording. (**b**) Post-hoc verification of the electrode tracks and optic fiber placement. (**c**) Post-hoc verification of viral expression in the EC (left) and in CCK-positive projections in the LA (middle). A magnified image is shown in the right panel and corresponds to the boxed area of the middle panel. (**d**) Co-immunofluorescent staining of the CCK-positive fiber (EYFP), the axon terminal (synaptophysin), and CCKBR in the LA. The white arrowhead indicates a triple-positive neural terminal. Quantification of the CCK and CCKBR double-positive neural terminals out of all CCK-positive terminals (right). (**e**) Representative traces of auditory evoked potential (AEP) and visual evoked potential (VEP) at different sound and light intensities. (**f**) AEP and VEP peak latency. (**g**) Representative input/ouput (IO) curves for AEP (left) and VEP (right). (**h**) Detailed pairing protocol to induce LTP. Representative averaged fEPSP trace evoked by HFLS is shown in the inset. HFLS, high frequency laser stimulation; AS, auditory stimulation. (**i**) Time course plot of the normalized slope of AEP and VEP during LTP. The arrow indicates the application of LTP induction. Representative traces of averaged AEP/VEP before (–10–0 min, dotted line) and after (10–20 min, solid line) induction from the three groups are shown in the top insets. (**j**) The average normalized slopes 10 min before pairing (–10–0 min, before) and 10 min after pairing (10–20 min, after) in the three groups. \*\*\**P* < 0.001; two-way RM ANOVA with Bonferroni post-hoc pairwise test; RM ANOVA, repeated measures analysis of variance; NS, not significant.

In the next experiment, we examined the possibility of other neuroactive molecules that are co-released with CCK and contribute to HFLS-induced AEP-LTP. We adopted an RNA interference technique that specifically knockdown the CCK expression in the EC. We accomplished this by injecting a Cre-dependent AAV cassette carrying a ChR2 variant (E123T/T159C) and a short hairpin RNA (shRNA) targeting CCK (anti-CCK) or a nonsense sequence (anti-Scramble) into the EC of CCK-Cre mice (Figure 7a–c). The inclusion of laser-responsive ChR2 allowed us to induce the above AEP-LTP by specifically stimulating the EC^CCK+^→ LA pathway. We applied our HFLS pairing protocol in these mice and found that AEP-LTP could not be induced in the anti-CCK group but could successfully induced in the anti-Scramble group (Figure 7d–f, two-way RM ANOVA, significant interaction, F [1,31] = 14.94, *P* < 0.001; pairwise comparison, before vs. after pairing in the anti-CCK group, 101.5% ± 2.5% vs. 98.0% ± 4.8%, *P* > 0.05; before vs. after pairing in the anti-Scramble group, 103.0% ± 3.8% vs. 138.8% ± 9.7%, *P* < 0.001). This observation implies that CCK alone is responsible for HFLS-induced AEP-LTP.

**Figure 7.**
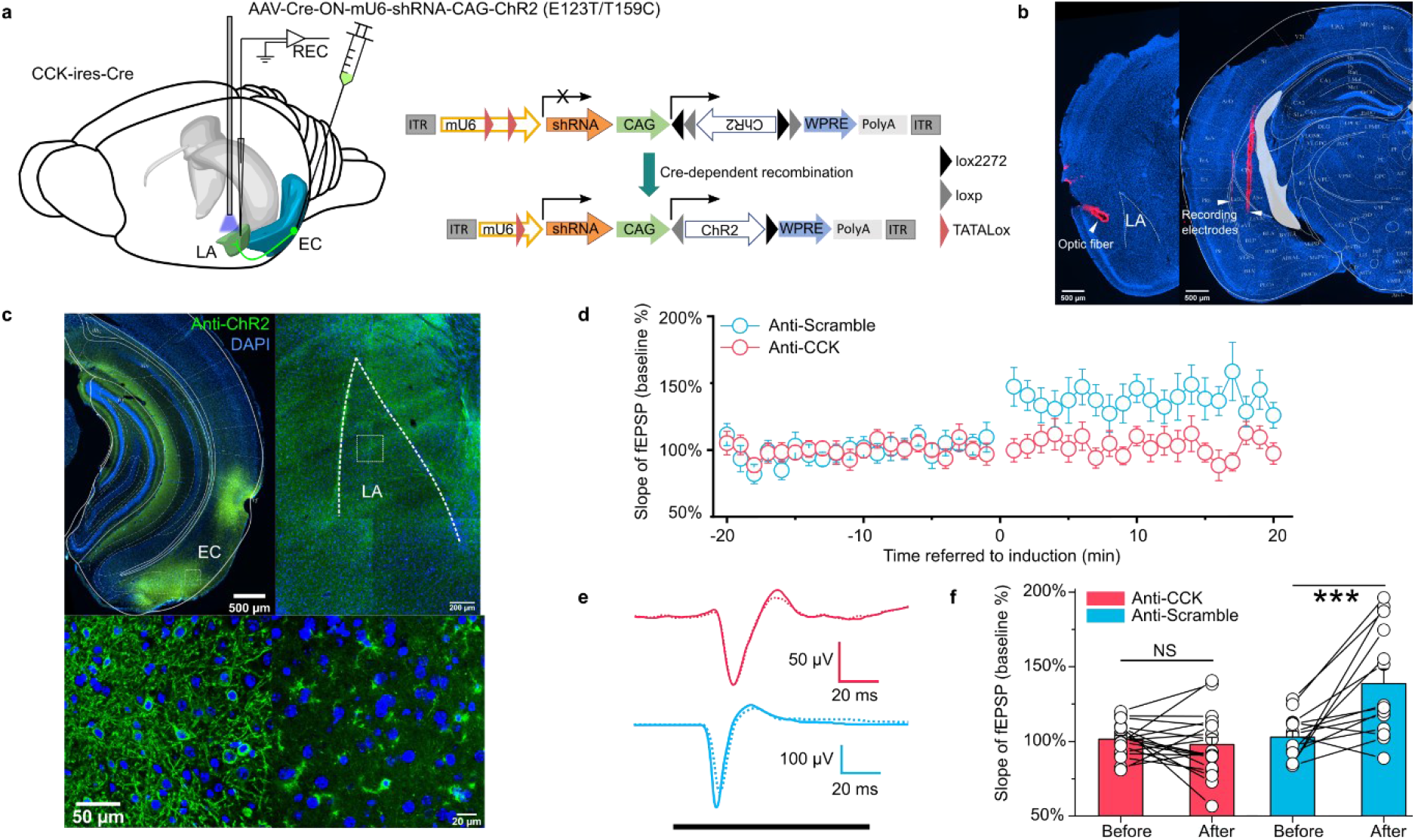
*In vivo* knockdown of CCK expression blocks AEP-LTP induction in the LA. (**a**) Schematic diagram of the experiment. CCK-Cre mice were injected in the EC with an AAV expressing shRNA (anti-CCK or anti-Scramble) and ChR2. *In vivo* recording was conducted in the LA (left). After Cre-mediated recombination, EC-CCK neurons were transfected with shRNA targeting CCK (anti-CCK) or nonsense sequence (anti-Scramble) as well as the excitatory opsin ChR2 variant E123T/T159C (right). AAV, adeno-associated virus; EC, entorhinal cortex; LA, lateral amygdala; REC, recording; ITR, inverted terminal repeat; mU6, mouse U6 promoter; CAG, CMV enhancer, chicken β-actin promoter; WPRE, woodchuck hepatitis virus (WHP) posttranscriptional regulatory element. (**b**) Post-hoc verification of the electrode tracks and optic fiber. (**c**) Post-hoc immunofluorescent staining targeting ChR2 in the EC (left) as well as in the CCK-positive projections distributed in the LA (right). Magnified images are shown in the bottom insets. (**d**) Time course plot of the normalized AEP slope before and after pairing in mice expressing anti-CCK or anti-Scramble shRNA. (**e**) Representative traces of the averaged AEP before (–10–0 min, dotted line) and after (10–20 min, solid line) induction in the two groups. Anti-Scramble is indicated in blue, and anti-CCK is indicated in red. (**f**) The average normalized slopes 10 min before pairing (–10–0 min, before) and 10 min after pairing (10–20 min, after) in the two groups. \*\*\**P* < 0.001, two-way RM ANOVA with Bonferroni post-hoc pairwise test; RM ANOVA, repeated measures analysis of variance; NS, not significant; fEPSP, field excitatory postsynaptic potential.

To dissect the real-time behavioral dependency of trace fear memory formation on the EC^CCK+^→ LA pathway, we employed optogenetics. We expressed the inhibitory opsin eNpHR3.0 (AAV-EF1α-DIO-eNpHR3.0-mCherry) or GFP control (AAV-hSyn-FLEX-GFP) in the EC of CCK-Cre mice. We also implanted optic fibers targeting the LA in these mice and then subjected the mice to trace fear conditioning (Figure 8a–b). During trace fear conditioning, EC^CCK+^→ LA were stimulated at a frequency of 5 Hz (i.e., 100 ms illumination + 100 ms interval) by the optic fibers for the duration of the CS and trace interval, as indicated in Figure 8a. For these experiments, mice were positioned in a head-fixed setup on a moveable surface, and an electrical tail shock was given as the US. After administration of the US, we most commonly observed flight (running). Interestingly, we found that after a few training trials, some GFP control mice (3/6 animals, data not shown) began running before the US was given, suggesting that GFP mice associate the CS with the US and make predictions in subsequent training trials (Movie S11). In contrast, we observe much fewer conditioned defensive responses in the eNpHR group throughout the training process (1/8 animals and 2/40 observed training trials, data not shown, Movie S12). Additionally, we recorded the freezing percentages in response to the CS before and after head-fixed fear conditioning (Figure 8c–d). We found that mice in the eNpHR group showed impaired freezing percentages post-training compared to mice in the GFP group (Figure 8d, two-way RM ANOVA, significant interaction, F [1,12] = 19.20, *P* < 0.001; pairwise comparison, GFP vs. eNpHR post-training, 39.0% ± 2.0% vs. 12.2% ± 4.8%, *P* < 0.001; Movie S13, S14). We did not observe any differences between the two groups at baseline (Figure 8d, pairwise comparison, GFP vs. eNpHR at baseline, 12.7% ± 3.4% vs. 12.2% ± 4.8%, *P* > 0.05) or prior to the CS (Figure 8c, two-way RM ANOVA, interaction not significant, F [1, 12] = 0.67, *P* = 0.43; pairwise comparison, GFP vs. eNpHR at baseline, 15.0% ± 2.8% vs. 8.0% ± 1.7%, *P* > 0.05; GFP vs. eNpHR post-training, 19.3% ± 3.8% vs. 17.8% ± 5.4%, *P* > 0.05). Altogether, our results suggest that trace fear memory formation is disturbed by real-time inhibition of the EC^CCK+^→ LA pathway.

**Figure 8.**
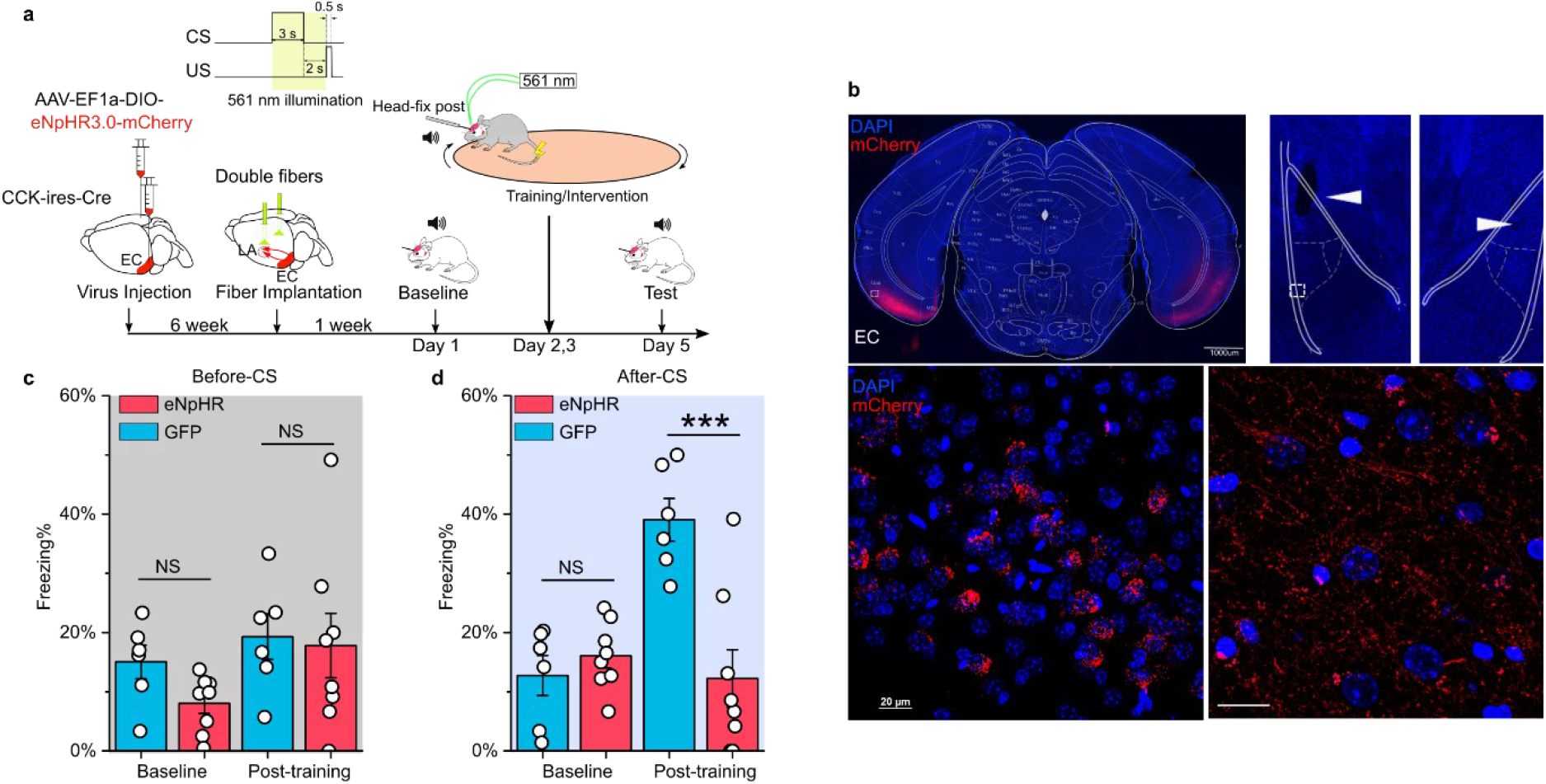
Real-time inhibition of the EC^CCK+^→ LA pathway impairs trace fear memory formation. (**a**) Schematic diagram of the experiment. The Cre-dependent inhibitory opsin eNpHR3.0 was expressed in the EC of CCK-Cre mice. Optic fibers were implanted near the LA to illuminate the CCK-positive fiber that signals from the EC to the LA during auditory-cued trace fear conditioning. The inset at the top right shows the timing of illumination, which covers the CS presentation and trace interval. EC, entorhinal cortex; LA, lateral amygdala; CS, conditioned stimulus; US, unconditioned stimulus. (**b**) Post-hoc verification of viral expression in the EC (top left) and of the optic fiber track in the LA (top right). The white rectangle in the top right panel is magnified in the bottom right panel. Magnified images show the transfected EC-CCK neurons (bottom left) and the CCK-positive EC-LA fibers (bottom right). (**c–d**) Freezing percentages before (**c**) and after (**d**) the CS in eNpHR-expressing mice (red, N = 8) and GFP-expressing control mice (blue, N = 6) on pre-training day (baseline) and post-training day. \*\*\**P* < 0.001; NS, not significant; two-way RM ANOVA with Bonferroni post-hoc pairwise test; RM ANOVA, repeated measures analysis of variance.

In summary, the release of the neuropeptide CCK from the EC neurons switches neural plasticity in the LA, and facilitates the formation of trace fear memory. Dysfunction in any part of this pathway impairs the formation of trace fear memory in mice. These results extend our understanding of learning and memory formation and have important implications for fear-related mental disorders.

## Discussion

Here, we employed classical Pavlovian trace fear conditioning to test the formation of trace fear memory in CCK^-/-^ and WT mice. We demonstrate that CCK^-/-^ mice have impaired fear responses compared to WT mice in both short and long trace fear conditioning. We also confirm that this behavioral defect is not caused by other abnormalities, including deficits in hearing and fear expression. Indeed, we demonstrate that depletion of CCK expression in mice impairs trace fear conditioning responses, which can be rescued by exogenous activation of CCKBR with its agonist CCK-4. Overall, our study suggests that trace fear memory formation and neural plasticity in the LA are dependent on a functional CCK network in the CNS.

Trace fear conditioning includes a gap between the CS and the US, which distinguishes it from the simultaneous CS-US termination in delay fear conditioning. In trace fear conditioning, mice must retain information from the CS during the trace interval and associate it with the subsequent US. As a result, the learning process in trace fear conditioning is slower than in delay fear conditioning, and fear generalization is more pronounced. We previously reported that WT animals form CS-US associations after three trials with minimal fear generalization in auditory-cued delay fear conditioning (X. Chen et al., 2019). In our previous report, we also demonstrated that CCK^-/-^ mice have difficulties in forming auditory-cued delay fear memory, visually-cued delay fear memory, or electrically-cued trace fear memory in which an electrical pulse stimulus in the auditory cortex is paired with a foot shock (X. Chen et al., 2019; Z. Zhang et al., 2020). Together, the results of our previous work and the present study indicate that the absence of the neuropeptide CCK has broad damaging effects on multiple forms of fear memory and is not limited to trace fear memory.

Fear conditioning can potentiate the signals of auditory responsive units in the LA (Quirk et al., 1995) in a phenomenon referred to as LTP. As a result, many studies have identified LTP as a physiological hallmark of fear conditioning (Blair HT, Schafe GE, Bauer EP, Rodrigues SM, 2001; Maren, 2001). In our study, we used *in vivo* recording to measure auditory-evoked field excitatory postsynaptic potential (fEPSP) or AEP. We did not find any apparent abnormalities in AEP (such as amplitude or latency) in CCK^-/-^ mice, suggesting that cortical and thalamic auditory inputs to the LA were functional. CCK^-/-^ mice did fail to induce AEP-LTP in the LA, strongly suggesting a deficiency in neural plasticity. However, we cannot simply assume that AEP-LTP induction is equivalent to trace fear memory. Occasionally, AEP-LTP is not sufficient to trigger the expression of fear behaviors. Kim and Cho reported that LTP in the LA was maintained during fear extinction (Kim & Cho, 2017). Thus, LTP in the LA is necessary but not sufficient for fear memory formation.

As the EC has been previously implicated in trace fear memory and behaviors, we manipulated EC function in our present study and investigated the behavioral and signaling outcomes. We found that silencing EC neurons with DREADD hM4Di impaired the formation of trace fear memory, which is consistent with several previous studies. For instance, electrolytic lesion of the EC impairs trace eyeblink conditioning performance in mice (Ryou et al., 2001), and neurotoxic lesions as well as M1 receptor blockade in the EC impair trace fear memory formation but not delay fear memory formation (Esclassan et al., 2009). Although this preferential association with trace fear memory has also been observed in certain areas of the hippocampus (Bangasser, 2006), the EC is a promising regulatory unit, because EC neurons maintain persistent spikes activity in response to stimuli (Egorov et al., 2002; Fransén et al., 2006). This sustained neuronal activity is thought to be the neural basis of ‘holding’ CS information during trace intervals to allow for CS-US association even after long trace intervals (20 seconds in our study). This information ‘holding’ theory is consistent with neuroimaging reports on working memory in subjects who ‘hold’ stimuli for specific periods (Nauer et al., 2015).

Auditory responses have been previously found in the EC and its upstream circuit (G. W. Zhang et al., 2018), however, these responses were limited to loud noise and did not involve the pure tone used in our behavioral paradigm. We reasoned that if the EC perceives and delivers the CS to downstream structures, then lesions in the EC would disturb the delay fear conditioning as well. Instead, previous studies have robustly demonstrated that EC lesions leave delay fear memory intact (Esclassan et al., 2009). The amygdala responds directly to the AS, and receives inputs from the AC, the MGB, and hippocampus directly. Thus, the EC is likely involved in the CS-US association in a more complicated manner, and this mechanism requires further investigation. We speculate that this mechanism is probably similarly as our previous finding in the sound-sound association (X. Chen et al., 2019) and visuo-auditory association (Z. Zhang et al., 2020), which is neuropeptide-based hetero-synaptic modulation machinery.

With cell type-specific tracing systems, we demonstrated that the EC is an upstream brain region that projects CCK-positive afferents to the LA, and these CCK-expressing EC neurons are primarily excitatory (Figure 3). Using anterograde Cre-dependent color switch labeling in the EC, we also found that CCK-expressing neurons were the predominant source of EC-LA projections, implying that CCK is integral to EC-LA connection and communication. Cell type-specific chemogenetic inhibition of CCK-expressing neurons in the EC also impaired the formation of trace fear memory. However, we cannot exclude the possibility that CCK may originate in other brain regions and contribute to fear memory formation.

We triggered the release of CCK from axon terminals after *in vivo* HFLS of CCK-expressing fibers in the LA (Hökfelt, 1991). In the presence of this artificially released CCK neuropeptide, we then presented the AS. The AS activates presynaptic axons via the canonical LA fear circuit, which is supported by the known role of the LA in receiving auditory input from both the auditory cortex and the thalamus (Romanski & LeDoux, 1992). In our study, the AS triggered postsynaptic neural firing. Therefore, our HFLS-mediated AEP-LTP induction protocol combines the released CCK with pre- and postsynaptic activation altogether in the LA and this pairing leads to the potentiation of AEP in the LA.

In the current study, we successfully excluded the contribution of substances co-released with CCK to the induction of AEP-LTP by applying the *in vivo* RNA interference to knockdown the expression of *Cck* in CCK-positive neurons of the EC. We found that knockdown of *Cck* blocked the induction of AEP-LTP and our *in vivo* application of shRNA supports the clinical use of shRNA to target mental disorders related to the CCK system. Our results that the inhibition of CCK-positive EC afferents to the LA impaired trace fear memory formation during both the learning and response phases suggest that establishing the CS-US association during trace fear conditioning requires functional CCK-positive EC-LA projections.

In conclusion, we found that EC-LA projections modulate neuroplasticity in the LA and therefore contribute to the formation of trace fear memory. The CCK terminals of the EC neurons in the LA release CCK that enable hetero-synaptic neuroplasticity of the auditory pathway to the LA. Our findings add a novel insight into the participation of the neuropeptide CCK in the formation of the trace fear memory. As various mental disorders, including anxiety (Davis, 1992), depression (Shen et al., 2019; Siegle et al., 2007), and PTSD (Shin et al., 2006), are highly correlated with hyperactivation and dysfunction of the amygdala and the fear memory circuitry, our finding supports CCK and its receptors as potential new targets for future therapeutic applications in these disorders.

## Author Contributions

JH, HF and XC designed the experiments; HF conducted the electrophysiological and behavioral experiments in mice; JS designed and manufactured two AAVs; HF, WF collected the data of behavioral experiments; HF, WF collected and analyzed the anatomy data; JH, and HF wrote the manuscript.

## Materials and Methods

### Animals

Adult male and female C57BL/6, CCK^-/-^ (CCK-CreER), and CCK-Cre (CCK-ires-Cre) mice were used in experiments. For behavioral experiments, only adult male mice were used. Mice were housed in a 12 hour light/12 hour dark cycle (dark from 08:00 to 20:00) and were provided food and water *ad libitum*. All experimental procedures were approved by the Animal Subjects Ethics Sub-Committee of the City University of Hong Kong.

For surgical procedures when doing virus injection and optic fiber implantation, mice were anesthetized with pentobarbital sodium (80 mg/kg, i.p., 20% Dorminal, Alfasan International B.V., Woerden, Netherlands,). For acute electrophysiological recording, mice were anesthetized with pentobarbital sodium (80 mg/kg, i.p.) or urethane sodium (2 g/kg, i.p., Sigma-Aldrich, St. Louis, MO, USA). Both anesthetics were periodically supplemented during the experiment to maintain anesthesia. Mice were fixed in a stereotaxic device, and the scalp was incised. A local anesthetic (xylocaine, 2%) was applied to the incision site for analgesia. After skull levelling, craniotomies were performed with varying parameters based on the region of the brain being accessed.

### Auditory and visual stimuli

Auditory stimuli, including pure tones and white noise, were digitally generated by a specialized auditory processor (RZ6 from Tucker-Davis Technologies [TDT], Alachua, FL, USA). For behavioral experiments, auditory stimuli were delivered via a free-field magnetic speaker (MF-1, TDT) mounted 60cm above the animal. Sound intensity was adjusted by a condenser microphone (Center Technology, Taipei) to ∼70 dB when it reached the animal. For *in vivo* recording, auditory stimuli were delivered via a close-field speaker placed contralaterally to the recording side. The sound intensity that induced 50%–70% of the maximum response was selected. Visual stimuli were generated by a direct current (DC)-driven torch bulb via the analog voltage output of the TDT workstation. Light intensity was roughly quantified as the value of the trigger voltage. For *in vivo* recording, the light intensity that induced 50%–70% of the maximum response was selected.

### Auditory brainstem response recording

Mice were anesthetized with pentobarbital sodium (80 mg/kg, i.p.) and placed on a clean and warm blanket in a soundproof chamber. A free-field magnetic speaker (MF-1, TDT) was placed 10 cm away from the right ear of mice. Recording, reference and ground needle electrodes (Spes Medica, Genova, Italy) were subcutaneously inserted below the forehead, right ear and left ear, respectively. Auditory stimuli (wide spectrum clicks, 0.1 ms) were presented to the mouse with a decreasing level from 80 dB to 20 dB with an interval of 5 dB. For each level of click stimulus, total 512 times of presentation were given at a frequency of 21 Hz. ABR signals were collected via a specialized processor (RZ6, TDT) and digitalized with a bandpass filter from 100 Hz to 5 kHz. Stimuli generation and data processing was performed with software BioSigRZ (TDT).

### Trace fear conditioning

On pre-conditioning day, each mouse was placed into the testing context (acrylic box with white wallpaper measuring 25 cm × 25 cm × 25 cm) for habituation and baseline recording. After 3 min of habituation, a CS (2.7 kHz or 8.2 kHz pure tone, 70 dB SPL, 3 s for the short trace paradigm and 10 s for the long trace paradigm) was given three times within 20 min.

On conditioning day, the mouse was placed into the fear conditioning context (acrylic box with brown wallpaper measuring 18 cm wide × 18 cm long × 30 cm high and equipped with foot shock stainless steel grid floor). After 3 min of habituation, a CS-US pairing was given. In the short trace interval paradigm, an US (0.5 mA foot shock, 0.5 s) was given 2 s after a 3-s-long CS. Three trials were given on each training day, and the interval between trials was 10–15 min. Totally two training days were given. The mouse was kept in the fear conditioning context for a 10 min consolidation period after the last training trial. In the long trace interval paradigm, an US was given 20 s after a 10-s-long CS. Eight training trials were given each training day, and the interval between trials was 2–3 min. The mouse was kept in the fear conditioning context for a 5 min consolidation period after the last training trial. After training, each animal was kept in a temporary cage and returned to their home cage after all individuals finished training.

On post-conditioning day (test day), the mouse was placed into the testing context. After 3 min of habituation, a CS was presented to the animal twice with a 2 min-long interval between stimuli. Two min after the last trial, the animal was transferred to a temporary cage and returned to its home cage after all individuals in its cage finished testing.

All contexts were cleaned thoroughly with 75% ethanol after each individual session. All of the above procedures were conducted in a soundproof chamber, and all videos (baseline, training, and testing) were recorded with a webcam (Logitech C270) set in the ceiling of the chamber. Videos were analyzed with a custom program based on an open-source platform (Lopes et al., 2015) (https://bonsai-rx.org). Briefly, the centroid of the animal was extracted from the videos. By comparing the coordinates of the centroid frame by frame, we then calculated the distance moved between two frames. The instant velocity of the animal was calculated by dividing this distance by the time span between two adjacent frames. The freezing percentage was defined as the percentage of frames with an instant velocity lower than the threshold of all frames in an observed time window. We compared the output of this program to results observed by the naked eye. Finally, we selected 0.1 (pixel^2^/s) as the appropriate moving threshold to define freezing. Freezing score was defined as the binary value (0 or 1) of time frame with instant velocity higher (0, ‘not freezing’) or lower (1, ‘freezing’) than the threshold. For freezing score plot shown in Figure 1, 2 and 4, freezing scores from all test sessions were averaged per second for data visualization.

### Electrophysiological recording in the LA and EC

Mice were subjected to the surgical procedures describe above. Tracheotomy was conducted to facilitate breathing and to prevent asphyxia caused by tracheal secretions during the experiment. Craniotomy was performed 1.0–2.0 mm posterior and 3.0–4.0 mm lateral to the bregma to target the LA. Dura mater was partially opened using a metal hook made of a 29G syringe needle. Tungsten recording electrodes (0.5–3.0 MΩ, FHC, Bowdoin, ME USA) were slowly inserted into the LA (approximately 3.5 mm from the brain surface). For laser stimulation experiments, another craniotomy was performed at the temporal lobe (1.0-2.0 mm posterior to the bregma) to expose the lateral rhinal vein. One optic fiber (200 µm diameter, 0.22 NA, Thorlabs, Newton, NJ, USA) was inserted below the rhinal vein and forwarded till 1.0–1.5 mm from the surface. The angle of the optic fiber was approximately 75° from the vertical reference. Responses were recorded and passed to a pre-amplifier (PZ5, TDT) and an acquisition system (RZ5D, TDT). Signals were filtered for field potential or spikes with respective bandwidth ranges of 10–500 Hz and 1–5000 Hz. All recordings were stored using TDT software (OpenEx, TDT). The maximum sound intensity was defined as the intensity that elicited a saturated AEP. The AEP baseline was recorded with 50% of the maximum sound intensity at a 5 s intertrial interval (ITI) for 20 min. For high-frequency electrical stimulation (HFS) experiments, we used ∼ 70% of the maximum sound intensity and a 150 µA electrical stimulation current. For high-frequency laser stimulation (HFLS) experiments, we used > 10 mW laser power to ensure activation of transfected axons. After AEP-LTP induction, we recorded the AEP for another 20 min.

For recording in the EC, we applied the protocol from the Li I. Zhang laboratory (G. W. Zhang et al., 2018). Craniotomy was performed at the juncture of the temporal, occipital, and interparietal bones and exposed the caudal rhinal vein and the transverse sinus (Figure S3). Electrodes were inserted approximately 1 mm below the dura mater.

All field potential data were extracted and processed in the MATLAB program, and all single unit data were extracted from the TDT data tank to the Offline Sorter (Plexon) for spike sorting. Sorted data were forwarded to the Neuroexplorer (Plexon) for additional processing and visualization.

### Plasmid construction and AAV packaging

The sequence and cloning details of plasmid will be described elsewhere (Su et al., manuscript in preparation). In principle, we generated AAV vectors that allow Cre-controlled expression of shRNA and channelrhodopsin in neurons. For plasmid pAAV-Cre-ON-mU6-ShRNA-CAG-ChR2(E123T/T159C), shRNA was placed under the control of a mouse U6 (mU6) promoter inserted with a TATALox element (Ventura et al., 2004). CAG-DIO-ChR2(E123T/T159C) cassette was inserted following the mU6-TATAlox-ShRNA cassette.

In brief, the pAAV backbone was recovered after digesting pAAV-CAG-Flex-tdTomato (Addgene 28306) with NdeI and HindIII. Fragment 1 (pUC57-Cre-ON-mU6-shRNA) was acquired by digesting pUC57-Cre-ON-mU6(TATALox) with HpaI and XhoI and then ligating it with annealed oligos that targets the coding sequence of Cck mRNA (Anti-CCK) or nonsense sequence (Anti-Scramble). Fragment 2 was acquired by digesting pUC57-CAG-DIO-ChR2(E123T/T159C)-Flag) with XhoI and HindIII. Fragment 3 was acquired by digesting pUC57-CAG-DIO-mCherry-EYFP (inverted)) with EcoRI and HindIII. pAAV backbone, Fragment 1 and Fragment 2 was ligated to make pAAV-Cre-ON-mU6-ShRNA-CAG-DIO-ChR2 (E123T/T159C)-Flag. pAAV backbone, Fragment 1 without shRNA, Fragment 3 was ligated to make pAAV-CAG-DO-mCherry-DIO-EYFP. DNA templates and shRNA oligoes mentioned above were acquired from Addgene or synthesized from BGI (Shenzhen, China) and verified by sequencing.

For AAV packaging (Xiong et al., 2015), HEK293T cells were seeded into 5 dishes (15cm, poly-D-lysine coated) for 1 viral preparation one day before transfection. Standard medium (DMEM, +10% FBS and antibiotics) were used for HEK293T cells. For PEI transfection, mix 35 μg AAV8 helper plasmid, 35 μg AAV vector, 100 μg pHGTI-adenol, 510 μL of PEI (1 μg/mL, Sigma) with DMEM (without FBS or antibiotics) to final volume of 25 mL. Incubate this mixture at room temperature for 15 min. Meanwhile, replace the media in dishes with DMEM + 10% NuSerum (Bio-gene) + antibiotics (20 mL/plate). Then add 5 mL of transformation mix per plate. 24 hours after transfection, change the culture media to DMEM + antibiotics without Serum. 72 hours after transfection, culture medium was collected and filtered to get rid of cell pellets. Collected medium was stirred at 4 °C for 1.5 hours, meanwhile mixed with NaCl (final concentration of 0.4 M) and PEG8000 (final concentration of 8.5% w/v). Virus were precipitated by centrifugation at 7000 g for 10 min. Supernatant was discarded and 10 mL lysis buffer (150 mM NaCl, 20 mM Tris pH = 8.0) was added to re-suspend the virus pellet. Virus was then concentrated and purified via Iodixanol gradients (“Optiprep” Sigma D1556-250mL). Centrifuge the gradients for 90 min at 46,500 rpm at 16 °C. The virus in 40% fraction was harvested and mixed with PBS and then transferred to an Amacon 100K columns-UFC910008 to remove the Iodixanol. Purity and titer of virus were then assessed by SDS-PAGE and SYPRO ruby staining (S-12000, Life technologies, Carlsbad, CA, USA).

### Viral and tracer injection

Mice were subjected to the surgical procedures described above. For viral injection into the EC, the following rostral parameters were used: Anterior-Posterior (AP) = 3.25 mm, Medial-Lateral (ML) = 3.80 mm, Dorsal-Ventral (DV) = 3.60 mm from the surface, volume = 100 nL. Similarly, the following caudal parameters were used: AP = 4.25 mm, ML = 3.60 mm, DV = 2.60 mm from surface, volume = 200 nL. For injection of tracer or virus into the LA, we used the following parameters: AP = 1.70 mm, ML = 3.40 mm, DV = 3.70 mm from the surface, volume = 200 nL. Craniotomy was performed after skull levelling and partial opening of the dura mater using a syringe needle hook (29G). We used the Nanoliter2000 system (World Precision Instruments [WPI], Sarasota County, FL, USA) for all infusions. Viral or tracer infusions were slowly pumped into brain tissue trough a fine-tip glass pipette filled with silicon oil at a speed of no more than 50 nL/min. After infusion, the pipette was left in the injection site for an extra 5–10 min before slow withdrawal. After withdrawal of the pipette, the scalp was sutured, and a local anesthetic was applied. The animal was returned to its home cage after awaking. For axon stimulation (observation), the virus was expressed for at least 7 weeks, and for cell body stimulation (observation), the virus was expressed for at least 4 weeks. For CTB tracer labeling, we perfused animals after 7 days of viral expression.

### Optic fiber implantation

Mice were subjected to the surgical procedures described above. Craniotomy was performed bilaterally to target the LA using the coordinates described above. Optic fibers (optic cannulae) were gently inserted into the LA (50–100 µm above the target area) and fixed with dental cement (mega PRESS NV + JET X, megadental GmbH, Büdingen, Germany). For head fixation, a long screw was fixed to the skull with dental cement at a 45° angle from the vertical axis.

### Fiber photometry

The commercial 1-site Fiber Photometry System (Doric Lenses Inc, Quebec, Canada) coupled with the RZ5D processor (TDT, USA) was used in the current study. Excitation light at 470 nm and 405 nm was emitted from two fiber-coupled LEDs (M470F3 and M405FP1, Thorlabs) and sinusoidally modulated at 210 Hz and 330 Hz, respectively. The intensity of the excitation light was controlled by an LED driver (LEDD1B, Thorlabs) connected with the RZ5D processor via the software Synapse. Excitation light was delivered to the animal through a dichroic mirror embedded in single fluorescence MiniCube (Doric Lenses, Quebec, QC, Canada) in a fiber-optic patch cord (200 μm, 0.37 NA, Inper, Hangzhou, China). The intensity of the excitation light at the tip of the patch cord was adjusted to less than 30 μW to avoid photobleaching. The emission fluorescence was collected and transmitted through a bandpass filtered by the MiniCube. The fluorescent signal was then detected, amplified, and converted to an analog signal by the photoreceiver (Doric Lenses). Finally, the analog signal was digitalized by the RZ5D processor and analyzed using Synapse software at 1 kHz with a 5 Hz low-pass filter.

Optical fiber implantation and fiber photometry were used to visualize CCK activity in vivo via a fluorescent sensor. Briefly, the GPCR-activation-based CCK sensor (GRAB_CCK_, AAV-hSyn-CCK2.0) was developed by inserting a circular-permutated green fluorescent protein (cpEGFP) into the intracellular domain of CCKBR (Jing et al., 2019). Binding of CCKBR with its endogenous or exogenous ligand (CCK) induces a conformational change in cpEGFP and results in increased fluorescence intensity, which we measured by fiber photometry.

### Chemogenetic manipulation

Each animal (with DREADD virus injection) received CLZ (0.5 mg/kg, Sigma-Aldrich, dissolved with 0.1% DMSO) or vehicle (sterilized saline with 0.1% DMSO) by intraperitoneal injection. After injection, animals were kept in transfer cages for 30 min to allow the drug to penetrate the blood-brain-barrier (BBB) and bind to the DREADD receptor (Gomez et al., 2017). Animals were then placed in conditioning boxes for further training.

### Optogenetic manipulation

CCK-Cre mice were injected with AAV-EF1α-DIO-eNpHR3.0-mCherry or control AAV-hSyn-FLEX-GFP. After 7 weeks, animals received bilateral optic fiber implantation as described above. Mice were allowed a 1-week recovery period to adjust to the head-fix setup. Baseline freezing percentages were recorded in the testing context on the pre-conditioning day as described above. On the conditioning day, mice were head-fixed, and limbs were allowed to move freely on a smooth-rotatory round plate. Optic cables were connected to the implanted optic cannulae after cleaning the cannulae ends with 75% alcohol. Short trace training procedures were performed as described above with two exceptions. First, the US was delivered to the tail by attached wires. Second, the current was increased to 1.0 mA, because the fur on the tail can hamper perception of electrical shock. A 561 nm green laser (10–20 mW) was applied from the onset of the CS to the onset of the US with a frequency of 5 Hz (100 ms illumination + 100 ms interval). On post-conditioning day, the conditioned response of the animal was recorded in the fear conditioning context. All activity was captured by a camera on the ceiling and analyzed with the previously-described Bonsai program.

### Anatomy and immunohistochemistry

Animals were anesthetized with an overdose of pentobarbital sodium, perfused with ice-cold phosphate buffered saline (PBS, 0.01 M, Sigma-Aldrich), and fixed with paraformaldehyde solution (PFA, 4% in PBS, Santa Cruz Biotechnology, Dallas, TX, USA). Animals were decapitated, and the brain was gently removed and submerged into 4% PFA solution for additional fixation (∼48 hours). Brains were sectioned into 40-µm-thick slices on vibratome (Leica VT1000 S). To observe viral expression, neural tracer labeling, or electrode track verification, sections were counter-stained with DAPI (1:10000, Santa Cruz Biotechnology) for 10 min and mounted onto slides with 70% glycerol (Santa Cruz Biotechnology) in PBS. For immunohistochemistry, sections were washed with 0.01 M PBS three times for 7 min each and blocked with blocking solution (5% goat serum and 0.1% triton X-100 in PBS) at room temperature for 1.5 hours. Each primary antibody was diluted to the appropriate concentration (Table 1) in blocking solution and incubated on sections overnight at 4°C. The next day, sections were washed with PBS three times for 7 min each and stained with secondary antibody, which was prepared in PBST (0.1% triton X-100 in PBS). Each secondary antibody was incubated on sections at room temperature for 3 hours. After secondary incubation, the sections were washed with PBS three times for 7 min each and counter stained with DAPI for 10 min. Finally, sections were washed three times with PBS and mounted onto slides with 70% glycerol mounting medium. Fluorescent images were captured with a Nikon Eclipse Ni-E upright fluorescence microscope and a Zeiss LSM880 confocal microscope.

**Table 1.**
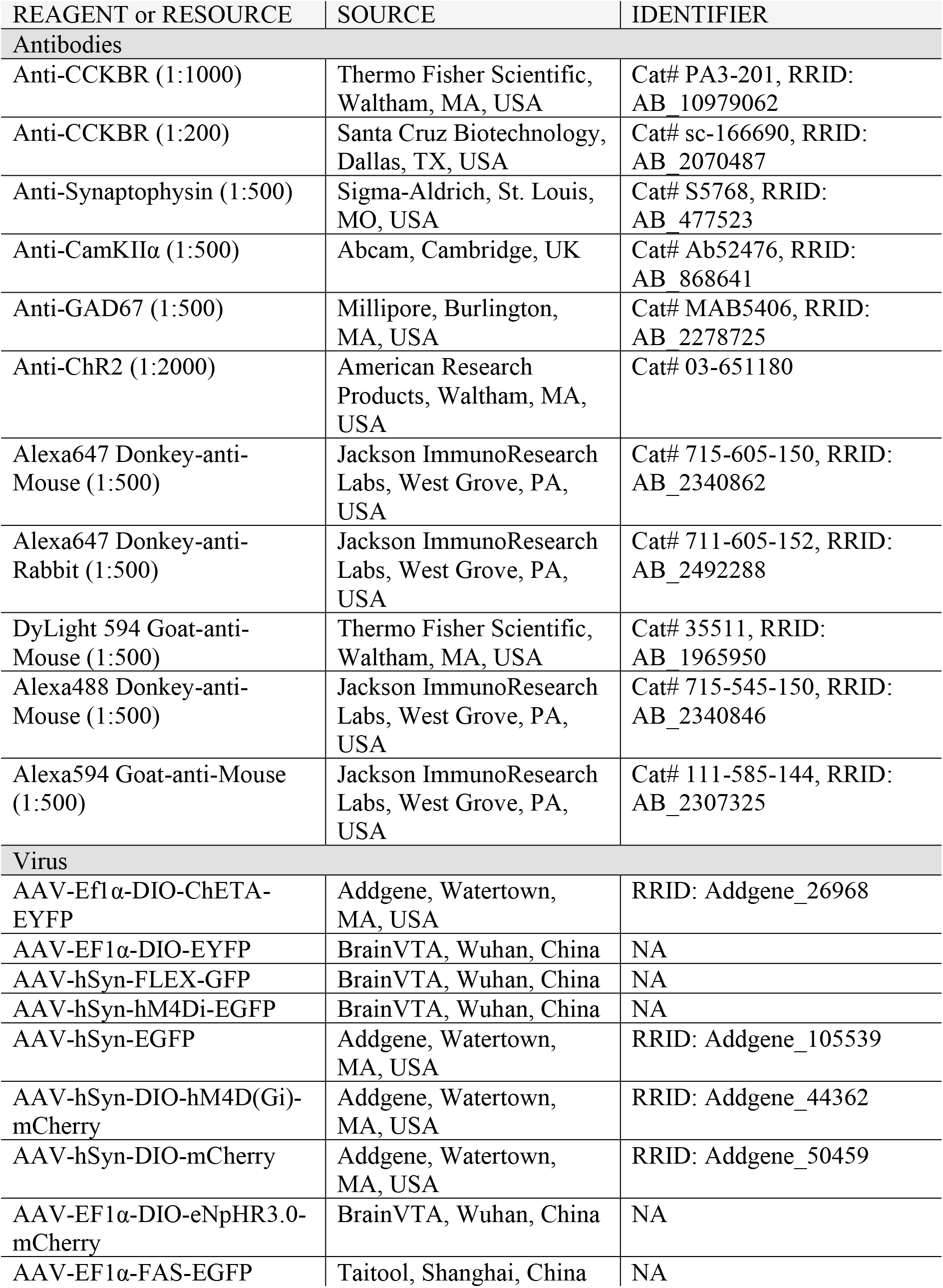
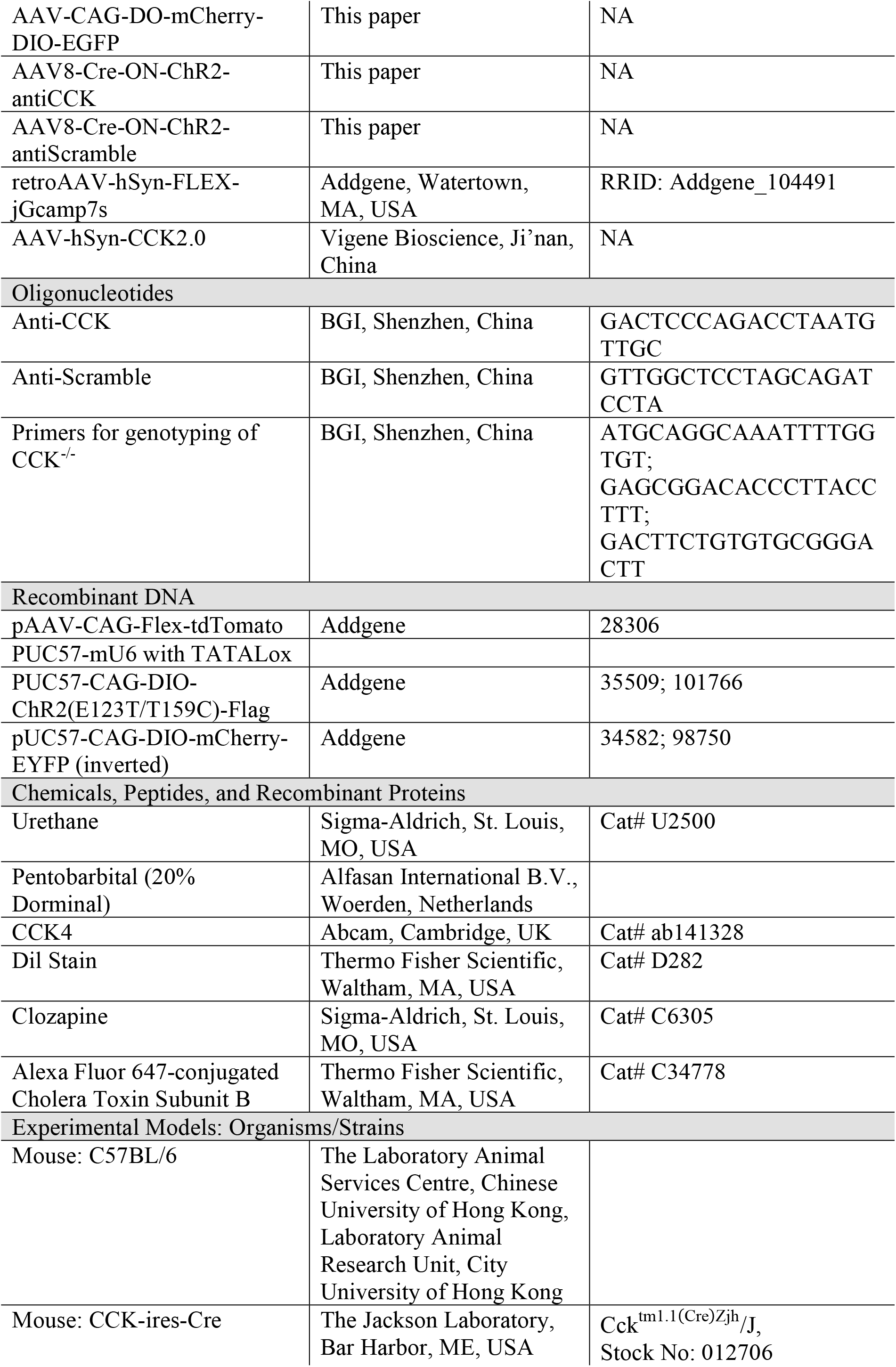
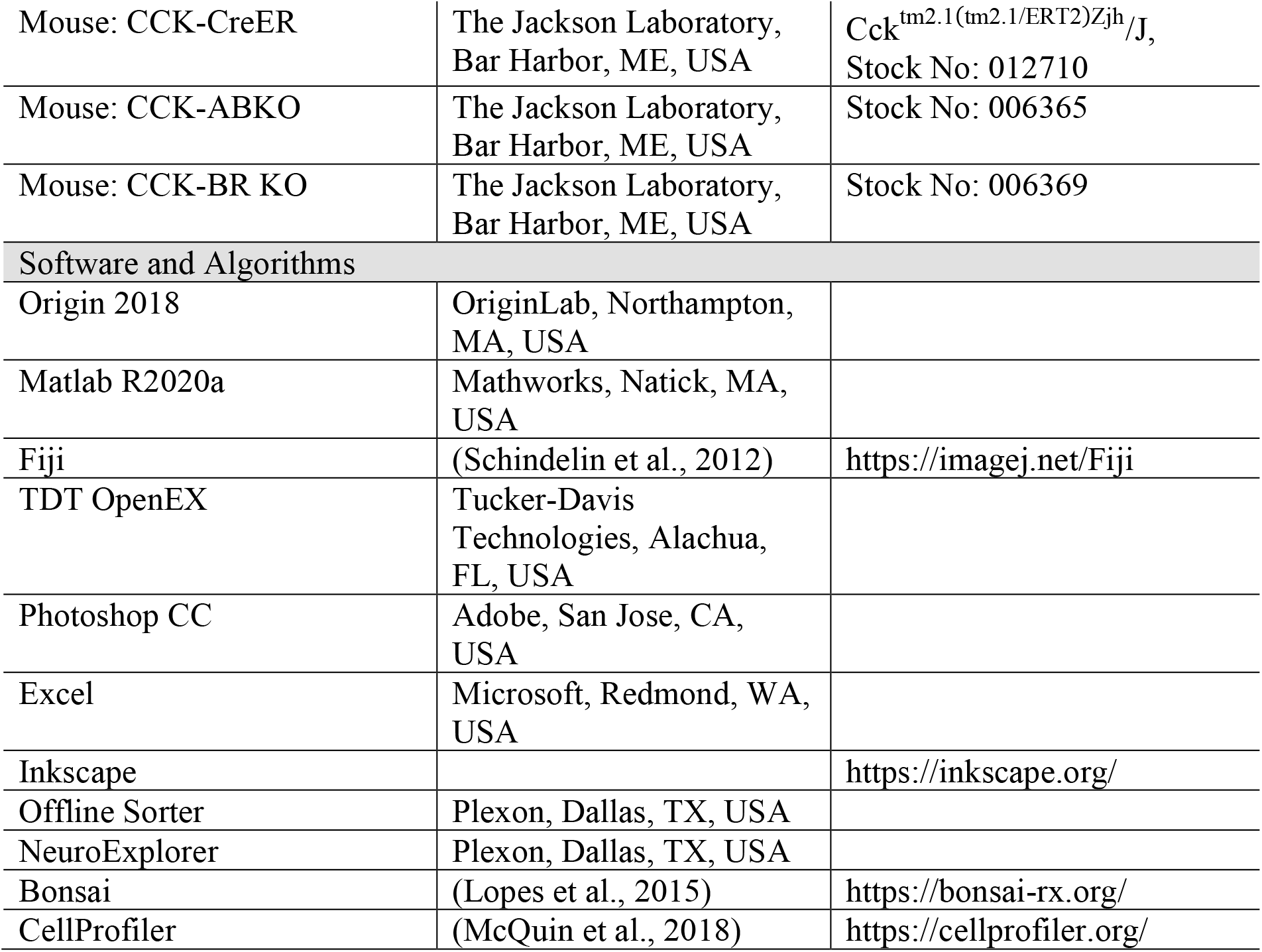
Key Resources.

### Image analysis

Imaging signal analysis, including quantification of intensity and percent positivity, was conducted in Fiji(https://imagej.net/Fiji) (Schindelin et al., 2012). To quantify the number (percentage) of viral- or immunohistochemical-positive neurons, we used the Cell Counter plugin in Fiji. To quantify the projection intensity of viral-positive neural fibers, we used the FeatureJ plugin in Fiji. We applied Hessian filter to extract the fiber-like structures and converted the raw images to eigen images with smallest eigen values selected. Eigen images were then converted to binary image by applying a threshold in Fiji and pixel density was measured as the intensity of neural projection (Grider et al., 2006). To quantify the colocalization of the CCK+ terminal (CCK-EYFP and synaptophysin double positive) and the CCKBR-innervating CCK+ terminal (CCK-EYFP, synaptophysin, and CCKBR triple positive), we extracted the double positive and triple positive pixels in Fiji and adopted the pixel-based colocalization analysis algorithm from CellProfiler (https://cellprofiler.org/examples) (McQuin et al., 2018) to calculate the colocalization ratios.

### Statistical analysis

Group data are shown as mean ± SEM (standard error of the mean) unless otherwise stated. Statistical analyses, including two sample t tests, paired sample t tests, one-way RM ANOVA (repeated measures analysis of variance), and two-way RM ANOVA, were conducted in Origin 2018 (OriginLab, Northampton, MA, USA). Statistical significance was defined as *P* < 0.05 by default.

## Supplementary Figures

**Supplementary Figure S1.**
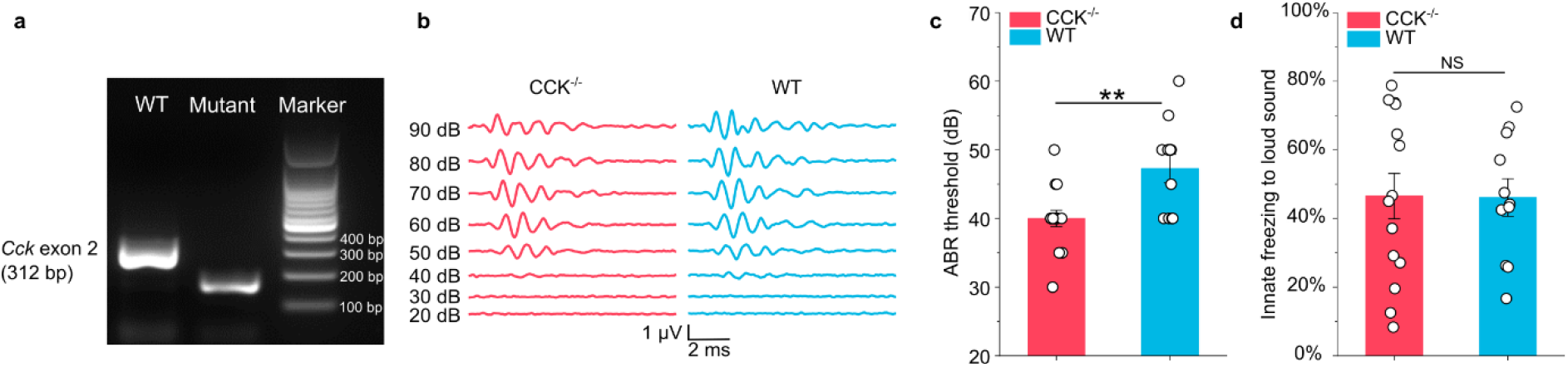
Genetic and behavioral examination of CCK^-/-^mice. (**a**) PCR-based genotyping results showing the absence of the *Cck* exon 2 (312 bp) fragment in the mutant sample. The band in the mutant sample is a fragment of the CreER gene. (**b**)Representative auditory brainstem response (ABR) traces from a CCK^-/-^ mouse and a WT mouse, respectively. (**c**) ABR thresholds in WT (N = 11) and CCK^-/-^ (N = 15) mice. ** *P* < 0.01; two-sample t-test. (**d**) Innate freezing levels in WT (N =14) and CCK^-/-^ (N = 10) mice. NS, not significant; *P* > 0.05; two-sample t-test.

**Supplementary Figure S2.**
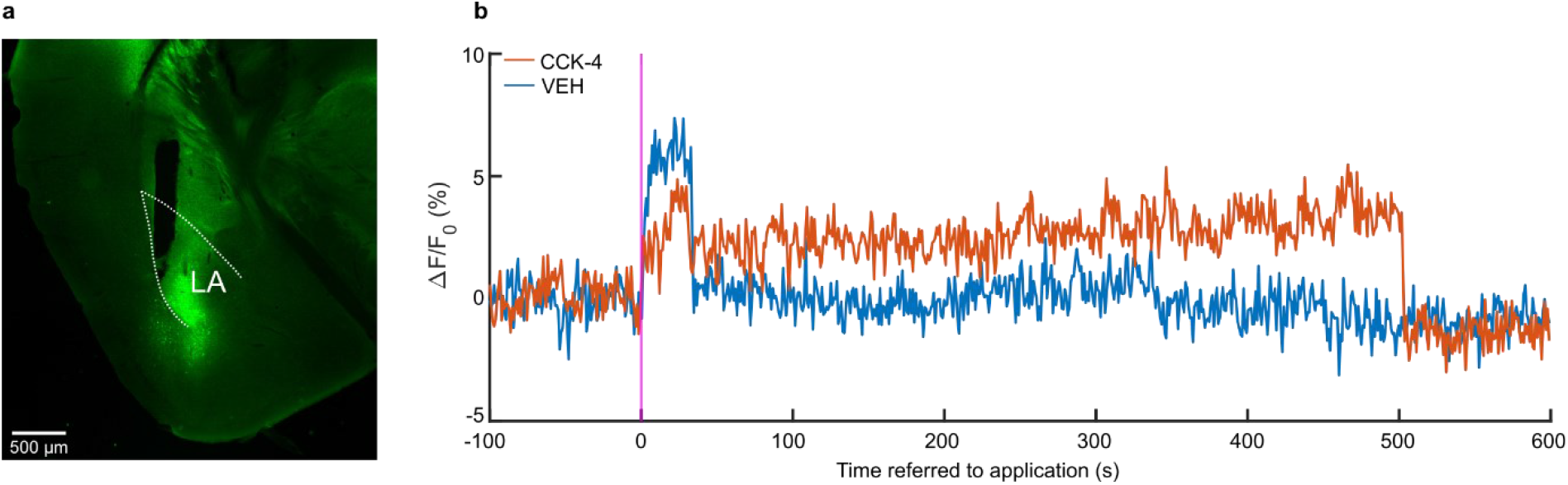
Exogenous CCK-4 activates CCKBR in the LA. (**a**) Verification of CCK-sensor2.0 expression and the optic fiber track in the LA. (**b**) Representative traces of fluorescence signal of the CCK-sensor before and after the peripheral application (intraperitoneal injection) of CCK-4 (1 mM, 200 μL) or vehicle (VEH). Fluorescence signal was measured by fiber-photometry with an implanted optical fiber in the LA.

**Supplementary Figure S3.**
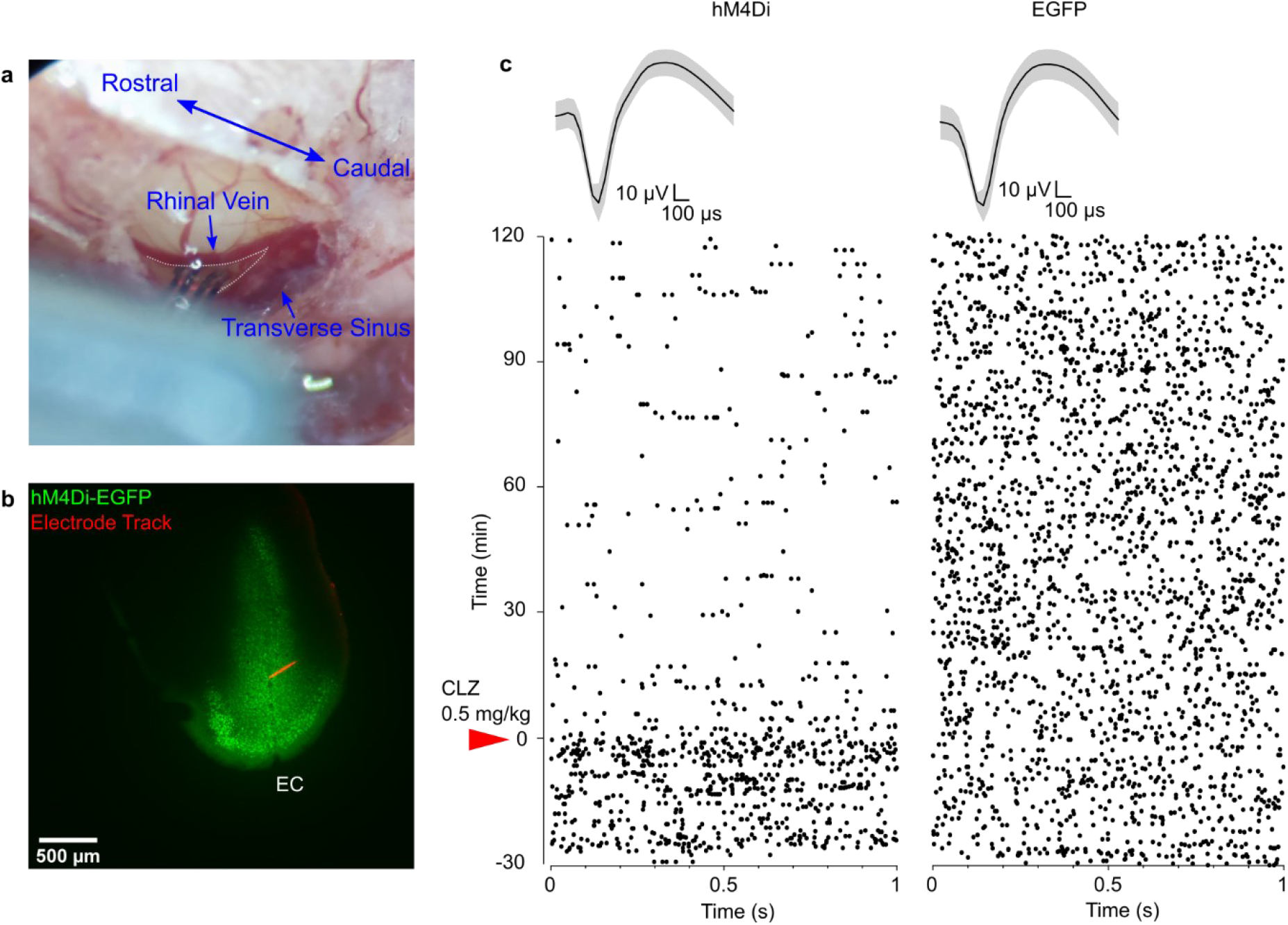
Verification of chemogenetic suppression in the EC via *in vivo* electrophysiological recording. (**a**) Image showing the location of the *in vivo* recording in the mouse. The caudal rhinal vein and the transverse sinus were as landmarks. The triangular area between these two veins (area defined by gray dotted line) was used to target the EC. (**b**) Post-hoc verification of hM4Di-EGFP viral expression and the electrode track, which was visualized using Alexa594-conjugated CTB. (**c**) Representative raster plots of single unit firing in the EC before and after intraperitoneal CLZ application in hM4Di-expressing (hM4Di, left) and EGFP-expressing (EGFP, right) mice. Waveforms of these two representative units are shown above the raster plots.

